# Bacterial colony biopsies: spatial discrimination of heterogeneous cell types by cytometric fingerprinting

**DOI:** 10.1101/2024.11.25.623575

**Authors:** Gorkhmaz Abbaszade, Kathrin Stückrath, Susann Müller

**Author notes:** Corresponding author Susann Müller, Department of Applied Microbial Ecology, Helmholtz-Centre for Environmental Research - UFZ, Permoserstr. 15, 04318 Leipzig, Germany.

## Abstract

Colonies of pure bacterial strains are highly dense cell structures that are organized in distinct and typical arrangements. The size, shape and variability of bacterial colonies are strongly dependent on the species and also influenced by environmental conditions. However, the spatial organization of individual cells is unknown for most strains. How specific heterogeneous cell types at different locations in a colony contribute to the overall structure and how they may influence the overall function of a colony is largely unknown. The study aims to investigate the local diversification of bacterial colony structures by introducing a local biopsy technique.

The biopsied cells were analyzed by microbial flow cytometry and cytometric fingerprinting, which diversified the biopsied samples into many heterogeneous cell states. This two-stage resolution insight into colony structure was performed on five bacterial strains: *Bacillus subtilis, Paenibacillus polymyxa, Kocuria rhizophila, Stenotrophomonas rhizophila*, and *Pseudomonas citronellolis*. The effects of biopsy tool size (27G needle and 10 µL, 200 µL, 1000 µL pipette tips) and sampling location on the precision of the technique were tested by using both gates setting along Gaussian distributions of subpopulations and a grid gating tool as well as the t-distributed Stochastic Neighbor Embedding (t-SNE) method.

The biopsy technique uncovered significant heterogeneity among the cells within bacterial colonies, identifying differences in cell cycle stages, the proportion of living and dead cells, and the abundance of spore types. Cells from different biopsy sites displayed distinct physiological states, revealing that colony structure is far more complex than previously understood. The technique’s precision depends on the biopsy tool size, dye equilibration, and cell handling, underscoring the importance of method calibration. The biopsy method, combined with cytometric fingerprinting, provided insights within only 15 to 45 minutes and is universally applicable.

The study provides a high-resolution biopsy technique that explores the spatial distribution of cell types and their heterogeneous physiological cell states, allowing conclusions to be drawn from biopsy composition at different locations to overarching functions of the entire bacterial colony. This method also facilitates downstream analysis through further cell sorting, offering a powerful approach for future functional investigations.

## INTRODUCTION

Microorganisms constitute the largest biomass on Earth (Bar-On et al., 2018). They predominantly exist in biofilms, a collective lifestyle distinct from individual planktonic cells. The ratio of bacteria living in biofilms to planktonic bacteria is estimated to be within a range of 10^2^ : 1 to 10^3^ : 1 (Magnabosco et al., 2018). Collective life forms offer several advantages over solitary existence, facilitating emergent functions that ultimately evolve from synergistic interactions that would not have arisen otherwise (Bingham & Ratcliff, 2024; Fisher et al., 2013; Flemming & Wuertz, 2019; Lyons & Kolter, 2015).

In a pure biofilm population, the nature and extent of emergent functions depend on the type, number, and distribution of encompassed heterogeneous cells and their physiological states, on the interaction between the phenotypes, as well as between cells and the environment (e.g., Liu et al., 2015). Collective life forms are prevalent because they provide fitness benefits and can mitigate conditions that might otherwise lead to extinction (Dalziel et al., 2021). Such relationships can be mediated through quorum sensing, gene regulation, or substrate sharing, and also through ecological paradigms such as dispersal within and between cell subsets, or through niche differentiation mechanisms that manifest the conditions for growth and survival.

To comprehensively understand the collective life of microbial populations and in particular the associated emergent functions, it is essential to study their spatial resolution in multiple dimensions (1D, 2D, or 3D) and ideally over time (4D) (Drescher et al., 2016; Hartmann et al., 2019; Yordanov et al., 2021). Three-dimensional structural insights are achieved through a variety of spectroscopic and microscopic techniques, which can provide information on physicochemical characteristics related to local population performance such as the synthesis of siderophores or extrapolymeric substances (Bien et al., 2022; Brockmann et al., 2019). Moreover, the structural organization of populations, the differentiation of cells into different phenotypes, and their development into multicellular entities can be studied at the single-cell level using confocal microscopy, transmission, and scanning electron microscopy or atomic force microscopy (see for an overview: Khan et al., 2020; Schlafer & Meyer, 2017). Additionally, 3D scalar mapping and 4D visualization algorithms are available for assessing and visualization of dynamic developments of biofilms (e.g., Hartmann et al., 2021; Paula et al., 2020).

Similar to confocal imaging techniques, flow cytometry (FCM) offers characterization of individual cells. This can be achieved through non-invasive methods such as light scattering, natural autofluorescence induced by cellular pigments, or a variety of intrinsic fluorescent reporters that indicate heterogeneous gene expression. In addition, large numbers of extrinsic fluorescent markers allow for specific labeling of physiological cell states. At first glance, FCM has the (surmountable) disadvantage of requiring single cells for analysis, which, in turn, requires disruption of the entire biofilm material. This challenge is outweighed by the major advantage of local and quantitative cell separation by fluorescence-activated cell sorting (FACS) that can be performed at high throughput (10^3^ to 10^4^ cells sec^-1^). The full potential of FCM-FACS analyses is realized when specifically highlighted (i.e., gated) bacterial cell types are sorted for further examination, such as through microscopy (Cichocki et al., 2020), cultivation (Bellais et al., 2022), gene expression studies (e.g., by mRNA-sequencing, Freiherr von Boeselager et al., 2018), or metabolic pathway analysis (e.g., by proteomics, Haange et al., 2020; Jehmlich et al., 2010). Since different cell types are distributed differently in colonies (e.g. Zhang et al., 2020), local biopsies offer the opportunity to study specific subpopulations and their heterogeneous cell types independently of the entire colony.

This study aims to understand the biogeographical structure of bacterial colonies by developing a standard procedure for extracting small biopsies from any part of a biofilm to identify and quantify local cell heterogeneity at the single-cell level. This required i) a local sampling of minimal numbers of bacterial cells, for which four different tip and needle sizes have been tested. The cell handling also required ii) the adaptation of existing flow cytometric cell treatment methods (Cichocki et al., 2020) to low cell numbers, including a DNA staining method to detect growth-related cell cycle and cell division states as well as spore types. In addition, the detection of dead cells relative to the number of vegetative cell types would allow, for example, the detection of suicidal cell-cell interactions as described for *B. subtilis* (Höfler et al., 2016). The new methods should also (iii) ensure cell handling that does not affect physiological cell states. Furthermore, for future functional identification, extracting and analyzing the mRNA content of individual bacterial cells presents unique challenges compared to eukaryotic cells (Longo et al., 2021). The total amount of mRNA in bacterial cells is two orders of magnitude less than in eukaryotic cells, and the half-life of mRNA is only a few minutes due to the absence of the 3’-polyadensosine tail (McNulty et al., 2023; Milo & Phillips, 2015). Significant efforts are currently being made to analyze the mRNA content both for sorted subpopulations (Freiherr von Boeselager et al., 2018) and for single cells (sc-mRNA, e.g. probe-based: McNulty et al., 2023; droplet-based BacDrop: Ma et al., 2023; MATQ-seq: Imdahl et al., 2020), using FACS in some applications. Consequently, the newly developed method should enable the potential future use of FACS for this purpose and iv) ensure rapid handling of cells. The effectiveness of the newly introduced methods should be v) universally applicable and therefore were tested on five bacterial strains, two gram-negative and three gram-positive, including two spore formers.

The proposed workflow will enable an unprecedented exploration of biofilm populations, facilitating the identification of specific and overarching functions and deciphering paradigms of population ecology at the scale of pure bacterial biofilms.

## MATERIALS AND METHODS

### STRAINS

For the biopsy experiments, five non-pathogenic bacterial strains, *Bacillus subtilis*, *Paenibacillus polymyxa*, *Kocuria rhizophila*, *Stenotrophomonas rhizophila,* and *Pseudomonas citronellolis*, were selected. They have well-characterized GC content and are fully sequenced. They exhibit different Gram-stain behavior, vary in genome sizes, and display diverse sporulation characteristics. Notably, *P. polymyxa*, *K. rhizophila,* and *S. rhizophila* are part of a standard cytometric mock community, as described by Cichocki et al. (2020) and Koch et al. (2013). Additionally, the strain *P. citronellolis* contains a fluorescent marker mScarlet (Table 1).

**Table 1:**
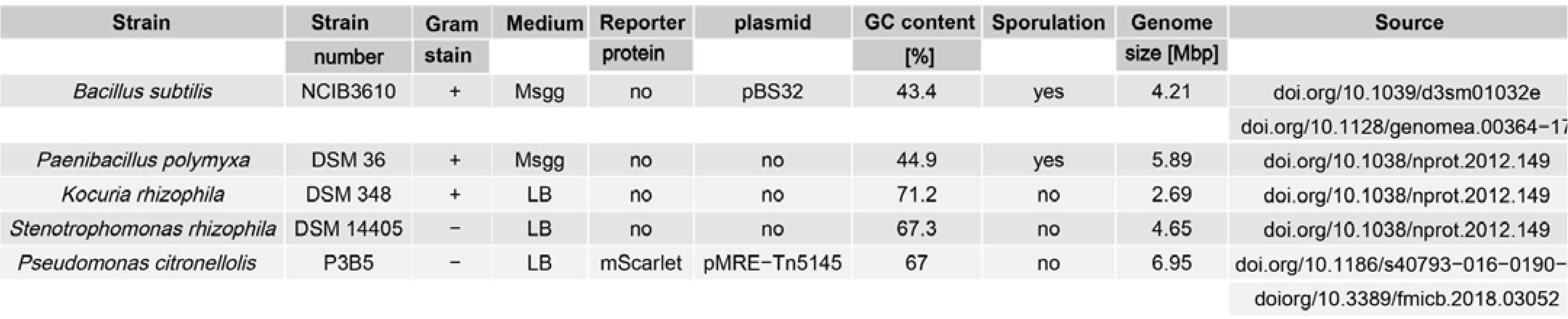
General features of bacterial strains, media used for growth, and literature for detailed information.

### CULTIVATION

All bacterial strains were pre-cultured in liquid culture. One colony of *B. subtilis* was inoculated in 20 mL DSMZ medium 1 for *Bacillus* and grown at 125 rpm and 30°C for 24 h. One colony from each of the strains *P. polymyxa*, *K. rhizophila*, *S. rhizophila*, and *P. citronellolis* was inoculated in 20 mL LB medium and grown at 125 rpm and 30°C for 24 h. Following, 2 µL of each of the preculture was either pipetted on a MSgg agar plate (*B. subtilis*; Branda et al., 2001; Freese et al., 1979) or on LB agar plate (*P. polymyxa, K. rhizophila*, *S. rhizophila*, *P. citronellolis*) at 30°C for 3 days, respectively.

### HARVESTING

For whole colony harvesting, a single colony was removed with a sterile spatula from the agar plate and resuspended in 2 mL PBS (6 mM Na_2_HPO_4_, 1.8 mM NaH_2_PO_4_, 145 mM NaCl; pH 7.2; 0.2 µm filtered). The cells were separated by swirling the solution 10 times using a 2 mL syringe with a 23G needle. After, the cell suspension was sonicated (20% duty cycle, 200 W) for 2 x 12 sec with Branson Ultrasonics^TM^Microtips and a Branson Sonifier 250 (Branson Ultrasonics^TM^, Brookfield, CO, USA).

The colony biopsy was performed using a manual punching machine (LYXM-STP-JYCBD FBA; Shenzhen Fast To Buy Co., Ltd.; Hong Kong, CN), which allows a sample to be taken from a bacterial colony at the same angle and line every time. The depths of biopsies were adjusted to the respective sample, different pipette tips and syringe needle size: 1000 µL tip (outer I] 1.3 mm; inner I] 0.83 mm), 200 µL tip (outer I] 0.9 mm; inner I] 0.52 mm), 10 µL tip (outer I] 0.75 mm; inner I] 0.35 mm), and 27G needle (outer ø 0.41 mm; inner ø 0.21 mm). The biopsy samples were resuspended in 50 µL PBS and rinsed by the tips or the 27G needle before 10 min sonication (35 kHz, 80 W; Ultrasonic bath DT 100 Sonorex Digitec; Bandelin, Berlin, Germany). Biopsies were obtained from all strains from the innermost to the outermost regions of the colony (SI Figure 1). Additional triplicate samples were specifically obtained from position 2 and 4, to ensure reliability and reproducibility of the data (Figure 1) with the best fitting needle or tip for taking a biopsy from every strain: 200 µL tip for *B. subtilis,* 10 µL tip for *P. polymyxa* and *P. citronellolis*, and the 27G needle for *K. rhizophila* and *S. rhizophila*.

**Figure 1:**
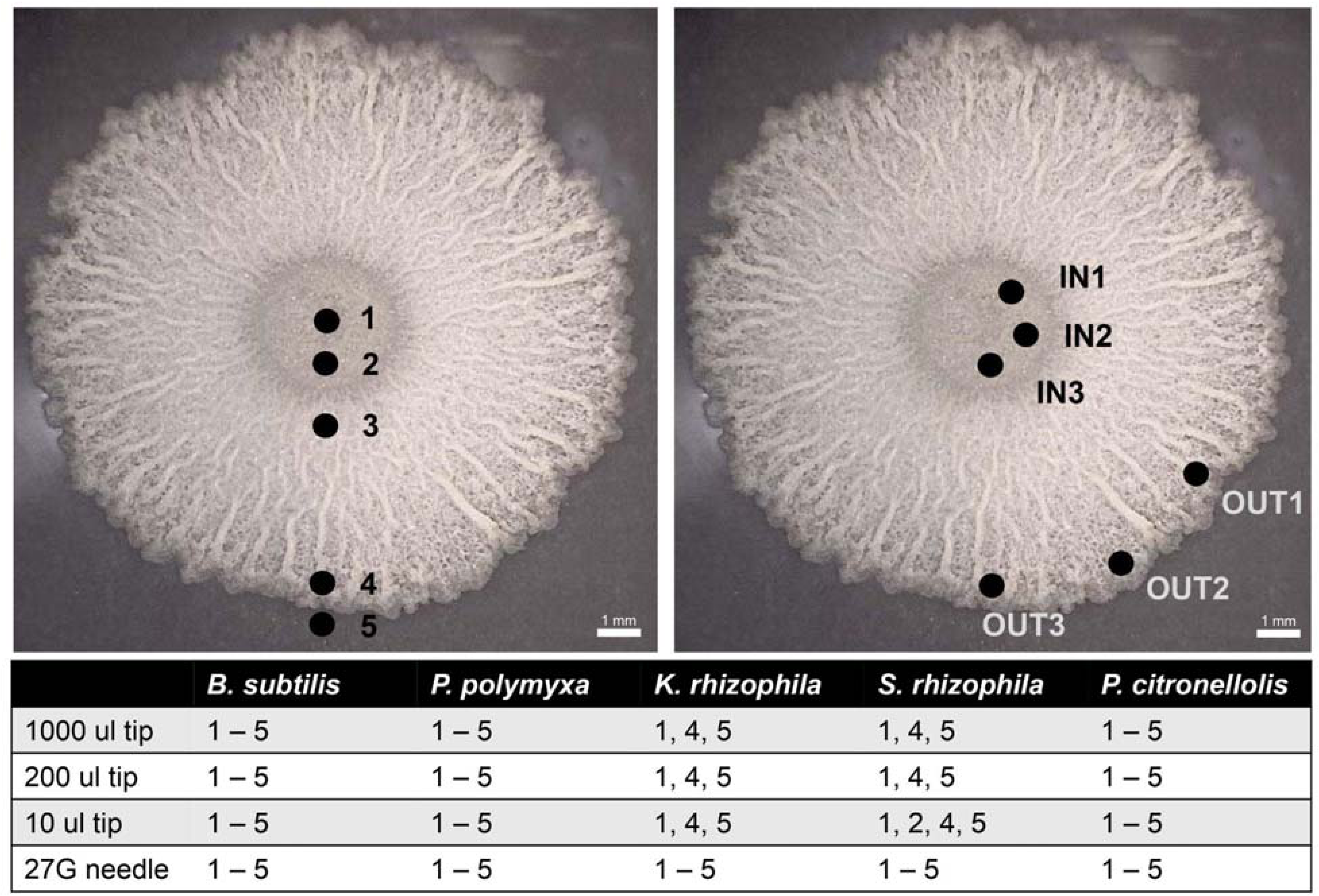
Image of a colony (here *B. subtilis*) and the sites where biopsies were taken from the colonies of all five strains. Left: biopsies taken from inside out. Right: Triplicates taken from position 2 and 4. Table 2: Number and location of biopsies taken from the five colony-grown strains using the different tip sizes and the 27G needle.

### CELL HANDLING

The cell density of samples from whole colonies was adjusted to an optical density (OD) of 0.04 at 700 nm (d = 0.5 cm) using phosphate-buffered saline (PBS; Spectrophotometer Ultraspec III, Amersham Biosciences Europe). Similarly, cells biopsied from the different sites of a colony were also adjusted to the same OD of 0.04 at 700 nm using PBS and the NanoDrop One (Thermo Fisher Scientific, Waltham, MA, USA), with the exception of all samples from position 5, all *B. subtilis* samples, the 10 µL sample of *P. polymyxa* and all 27G samples, due to their low cell count. For these samples, PBS was added up to a volume of 100 µL. For cell fixation 50 µL of a cell solution were incubated in 237.5 µL 20% NaCl (30% (m/v) stock solution; Merck KGaA, Darmstadt, Germany), 17.5 µL 1% NaN_3_ (20% (m/v) stock solution; Merck KGaA, Darmstadt, Germany) and 50 µL 10% C_2_H_5_OH (70% (v/v) stock solution; Chemsolute, Renningen, Germany) according to a method of López-Gálvez et al. (2023). The cells were fixated within 10 min at room temperature (RT).

A fingerprint-like pattern, to analyze population heterogeneity related to cell cycle states, was obtained by using 4’,6-di-amidino-2-phenyl-indole (DAPI; Merck KGaA - Sigma-Aldrich, Darmstadt, Germany). DAPI is a semi-permeable blue-fluorescent dye that intercalates to AT-rich regions of dsDNA. For the current application cells need to be fixated as described above, followed by the removal of the fixative after 10 min centrifugation (3,200 x g, 10 min, RT). To obtain a final concentration of 3.3 µM DAPI per sample, the cells were then resuspended in 50 µL PBS and 100 µL of a 5 µM DAPI phosphate buffer solution (0.4 M Na_2_HPO_4_, 0.4 M NaH_2_PO_4_ x H_2_O; pH 7; 0.2 µm filtered). The 5 µM DAPI phosphate buffer solution was freshly prepared from a 143 µM DAPI phosphate buffer stock solution which can be stored in the dark at RT for several days. After 10 min of incubation at RT, reference beads (1 μm Fluoresbrite® BB Carboxylate Microspheres; Ref. 17458, Lot. 659681; Polysciences, Inc., Warrington, PA, USA) were added as an internal sample standard. After 45 min, including the fixation and cell staining steps, the samples are ready for measurement.

To distinguish between live and dead cells, a combination of SYTO9 (all cell green-fluorescent membrane permeant nucleic-acid stain; Ref. S-34854; Lot. 2397781; ThermoFisher Scientific, Waltham, MA, USA) and propidium iodide (PI, indicator for compromised cell membranes and cell walls; Ref. P4170; Lot. WXBD4453V; Merck KGaA - Sigma-Aldrich, Darmstadt, Germany) was applied to live cells. A working solution of 10 µM SYTO9 (in 0.85% NaCl; 0.2 µm filtered) was freshly prepared each day from a 5 mM stock solution and stored on ice. To achieve a final concentration of 0.5 µM SYTO9 per sample, 7.9 µL of the SYTO9 working solution was added to 150 µL of adjusted cell suspension. In addition, PI (105 µM stock solution in 0.85% NaCl) was added at a concentration of 1.8 μM per sample. During staining, samples were kept for 12 min at RT. Following this, 3 µL reference beads were added for biopsy sites 1 - 4 and 1 µL reference beads for biopsy site 5 to keep the cell-bead ratio (1 μm yellow-green (YG) fluorescent Fluospheres; Ref. F-8823, Lot. 2221782; ThermoFischer Scientific, Waltham, MA, USA) as internal standard. The entire procedure was completed within 15 min.

### CYTOMETRIC MEASUREMENT

For cytometric analysis a BD Influx v7 Cell Sorter, (Becton Dickinson, Franklin Lakes, NJ, USA) equipped with a stream-in-air nozzle of 70 µm was used. The blue 488 nm Sapphire OPS laser (400 mW) was used for measurement of the forward scatter (FSC, 488/10; PMT1; related to cell size), the side scatter (SSC, trigger signal, 488/10; PMT2; related to cell density), the SYTO9 fluorescence (530/40; PMT3), and the PI fluorescence (616/23; PMT5). The UV laser 355 nm Genesis OPS laser (100 mW) was used to measure DAPI fluorescence (460/50; PMT9). The 561 nm Cobolt Jive laser (150 mW) was used to measure the mScarlet fluorescence of *P. citronellolis* (590/20; PMT 15 and 615/24; PMT 16). The light was detected by Hamamatsu R3896 PMTs in C6270 sockets (Hamamatsu, 211 Hamamatsu City, Japan). The fluidics ran at 33 psi through a 70 μm nozzle and cells were detected equivalent to an event rate of 2,500 to 3,000 events sec^−1^. FACSFlow Sheath Fluid (Ref. 342003; Becton Dickinson, Franklin Lakes, NJ, USA) was used as sheath buffer. The calibration of the cytometer was performed in the linear range by using 1 μm blue fluorescent FluoSpheres (Ref. F-8815, Lot. 69A1-1; Molecular Probes, Eugene, OR, USA), 2 μm yellow-green fluorescent FluoSpheres (Ref. F-8827, Lot. 1717426; ThermoFisher Scientific, Waltham, MA, USA), and 1 µm crimson fluorescenct FluoSpheres (Ref. F8816, Lot. 24005W; ThermoFisher Scientific, Waltham, MA, USA). All samples were measured in logarithmic scaled 2D plots and stored in the Zenodo database under the following doi: https://doi.org/10.5281/zenodo.13791365.

### DATA ANALYSIS

To differentiate the cells from instrumental and background noise, as well as beads, a cell gate was set according to the recommendations of Cichocki et al. (2020). The gate templates for all strains and for both FSC vs. DAPI and SYTO9 vs. PI are documented in SI Figures 2-11. These gating strategies were used to differentiate the various subsets of cells and to quantify their respective relative cell abundances in 2D plots. The program FlowJo 10.0.8.r1 (FlowJo, Becton Dickinson, Franklin Lakes, NJ, USA) was used to directly analyze the raw .fcs files.

**Figure 2:**
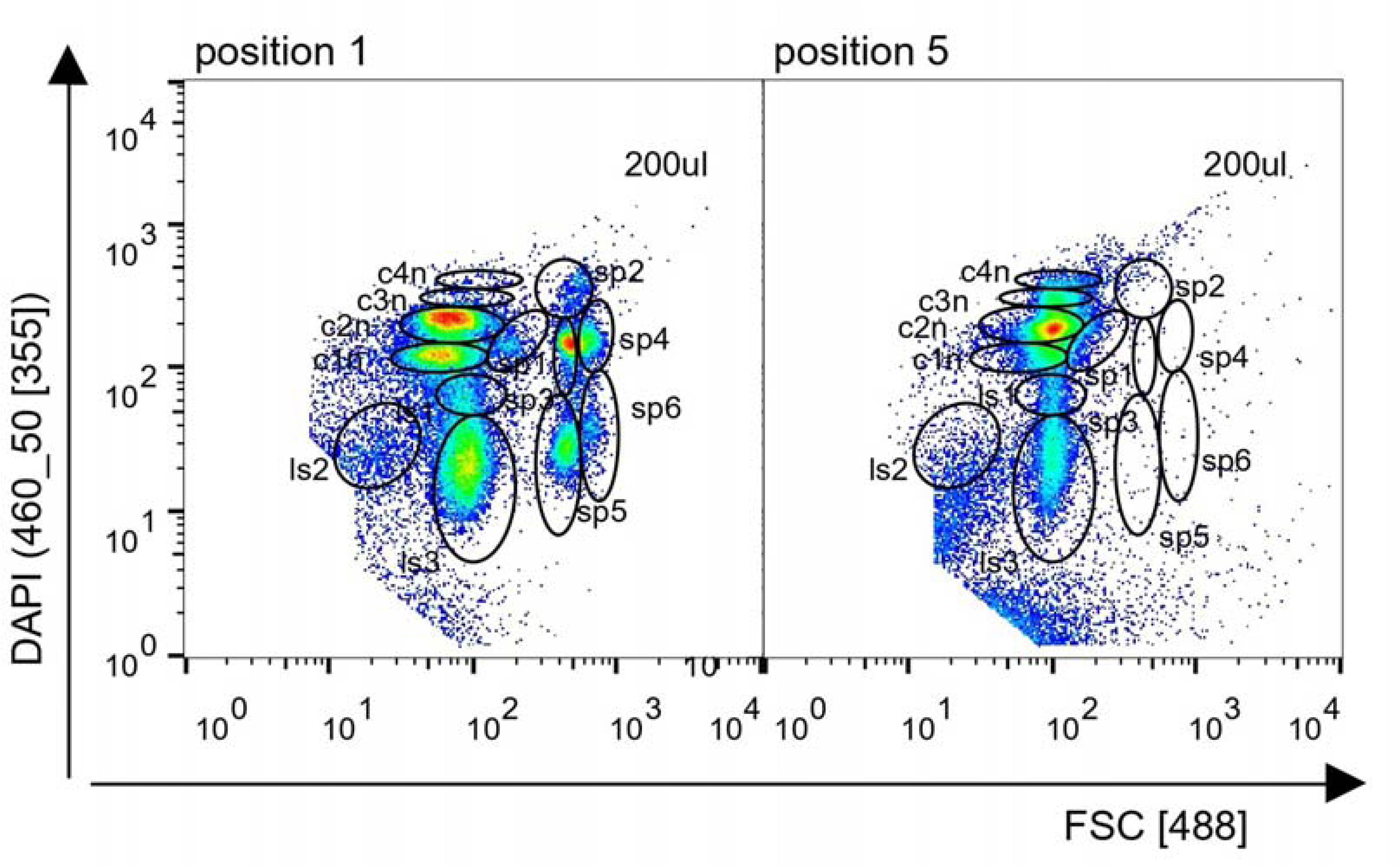
Cytometric FSC vs DAPI 2D plots of biopsy sites 1 and 5 of a B. subtilis colony using the 200 µL tip. The 2D FSC vs. DAPI 2D plots reveal 13 subpopulations (7 vegetative cell types and 6 spore types) at site 1 and 7 subpopulations at site 5 (7 vegetative cell types).

Subsequent data analysis incorporated the t-distributed Stochastic Neighbor Embedding (t-SNE) method for non-linear dimensionality reduction of the data (Hinton & Roweis, 2002). Before this, the cell counts were normalized using a Box-Cox transformation (Atkinson et al., 2021; Box & Cox, 1964) to ensure accurate comparative metrics.

To compare the in and out triplicate samples from five strains, we designed a gating strategy that divides the flow cytometry data into a 64-grid layout (8x8 gates). This template organizes the data into a matrix which allows to uniformly apply the gates across all samples. After collecting the corresponding cell numbers from all the gates, the cell counts from all strains were then merged to create a unified dataset. This combined data was subsequently used for dimensionality reduction t-SNE analysis, allowing us to visualize and compare the different cell types across the analyzed five strains in a reduced dimensional space. The plots were visualized by Rtsne (Krijthe, 2015) and vegan packages (Oksanen et al., 2024) in R.

Further boxplots were generated to effectively illustrate the distribution of various cell categories— namely live, dead, spores, and less stained cells, thereby enhancing the interpretability of the data. This was accomplished utilizing the ggplot2 (Wickham, 2016) and dplyr packages (Wickham et al., 2023) within R.

## RESULTS

### BIOPSY OF BACTERIAL PURE COLONY POPULATIONS

Biopsies of bacterial colonies allow the extraction of a limited number of cells from any location within a colony. Some colonies exhibited no apparent structural differentiation, presenting a visually unstructured appearance, while others had a clear texture of valleys and hill formations. Additionally, the physical characteristics of colonies, ranging from dry and firm to highly viscous and slimy, influenced the efficiency and accuracy of cell sampling. As the size of the colonies can vary depending on the bacterial strain and duration of cultivation, the accuracy of standard biopsy samples from 3-day-old colonies was investigated. Due to the variability of the colonies and the search for the best fit, different needle and tip sizes had to be tested, including a 27G needle and 10 µL, 200 µL and 1000 µL tips. Biopsies taken with the fine needle (27G) allow for the precise collection of very specific cell types, however, this approach may miss other surrounding important cell types. Conversely, large-tip size sampling may be nonspecific if colonies are too slimy. Biopsies were taken from five different strains: *Bacillus subtilis, Paenibacillus polymyxa*, *Kocuria rhizophila, Stenotrophomonas rhizophila*, and *Pseudomonas citronellolis,* either from the middle of the colony outwards or along concentric circles, as shown in Figure 1. Position 1 corresponds to the center, while position 5 is at the outer edge of the colony, with intermediate positions placed in between. During local biopsy, the selected cell types are separated from all other cells in the colony. This reduces any background noise caused by unsampled cells and provides clear and unbiased information about the selected cells. The cells obtained by this precise biopsy procedure are subjected to flow cytometric analysis to measure and characterize the heterogeneity of cell types at different biopsy sites.

### FLOW CYTOMETRIC CHARACTERISATION OF BIOPSIED CELLS

When analyzing biopsied cells via flow cytometry, two critical considerations must be addressed. Firstly, the developed cell handling methods should be able to measure biopsy samples with high specificity and reproducibility. This is particularly important because these samples may contain varying cell quantities, from very small to large numbers that could disturb the staining equilibrium. To this end, it is essential to identify the most appropriate tip or needle size for each bacterial strain to achieve both high local accuracy and a minimum number of cells required for subsequent individual cell characterization. Secondly, heterogeneous cell types from each biopsy can be sorted and further characterized in terms of their functional properties using downstream analyses. While this approach is well-established for proteomic studies (Haange et al., 2020), it has not been widely applied to transcriptomic studies. For potential subsequent transcriptomic analyses, rapid staining methods are required that can effectively differentiate cell heterogeneities.

To obtain information on heterogeneity of populations, a specific DNA staining method allows the number of chromosomes per cell to be determined (Müller, 2007), providing high-resolution information on cell cycle dynamics via FSC vs. DAPI 2D plot analysis. The heterogeneity of samples can be analyzed after 3 h using a standardized cell handling method (Cichocki et al., 2020) based on the amount of 3 x 10^8^ cells mL^-1^. In contrast, biopsy samples, especially those obtained using small tips or the 27G needle, contained far fewer cells. To overcome this limitation and reduce cell handling time, a 45-minute method (López-Gálvez et al., 2023) was adapted to colony-grown cells. This protocol was originally developed for online flow cytometry and did not include a centrifugation step. By comparing whole community samples treated with both the standard (PFA/C_2_H_5_OH fixation) and the 45-min (NaN_3_/C_2_H_5_OH fixation; SI Figure 12) procedures, the rapid applicability to colony-grown cells was verified. The new protocol revealed some additional, less stained (ls) subpopulations (e.g. ls2 and ls3 for *B. subtilis* and ls1 and ls2 for *P. citronellolis*). Only in the latter strain, these subpopulations were more abundant, highlighting those cell types that could not reach equilibrium with the dye due to unknown cell states. In principle, however, the distributions and degree of heterogeneity were the same for both methods in the FSC vs. DAPI 2D plots.

By using this method two degrees of diversification can be obtained from a colony: the location of the biopsy and the heterogeneity of cell types per biopsy. As an example, *B. subtilis* showed highly resolved cell cycle patterns and local biogeographic variations in cell heterogeneity for both biopsy positions 1 (13 subpopulations) and 5 (7 subpopulations; Figure 2). An overview of all biopsy sites analyzed for FSC vs. DAPI fingerprinting of *B. subtilis* is provided in SI Figure 2. The observed cell cycle patterns clearly indicated differential proliferation activities across various colony locations. The vegetative cells of *B. subtilis*, as shown in Figure 2 and SI Figure 2, underwent cell cycling by multiple replication of their chromosome numbers, as indicated by the position of the subpopulations in the 2D plot from c1n (100, relative fluorescence intensity, [rel. FI], y-scale) to c2n (200 [rel. FI]), to c3n (300 [rel. FI]), and to c4n (400 [rel. FI]). In addition, higher proportions of cells with higher chromosome numbers were found in position 5 compared to position 1 (c3n; c4n, increase in percentage from ∼1% to 8%), indicating higher cell cycle activities in the outer region of the colony. Spores that showed a higher FSC signal compared to the vegetative cells due to their compact structures were on the right side of the vegetative cells on the 2D plots (sp1 – sp6) and showed very different DAPI intensities. DAPI does not bind quantitatively in spores due to different supercoiled states of DNA depending on the sporulation state. The presence of spores was highest at position 3 of the colony for *B. subtilis* with a spore count of 51% and was dropping in position 4 to about 7 % (Figure 3, data from the 200 µL tip). Nearly no spores were found outside of the colony (position 5; Figures 2 and 3).

**Figure 3:**
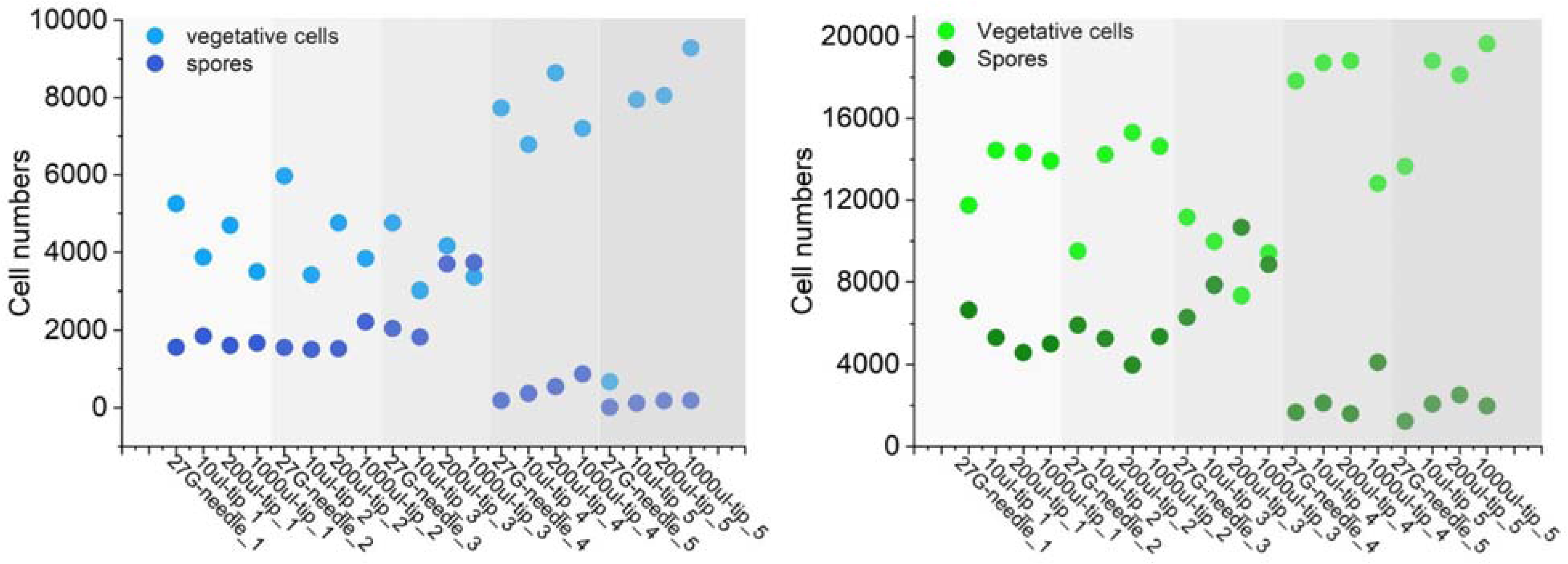
Distribution of abundances of vegetative cell types and spores of *B. subtilis*. The grey shades mark the positions of the biopsies taken with different needle and tip sizes. Left: DAPI-stained cell number. Right: SYOT9-stained cell number.

For the study of colony biogeography, the ratios between dead cells, vegetative cells, and spores are also crucial parameters in addition to cell cycle activities. To assess these ratios, biopsy specimens from the same sites were analyzed using the same tips and the 27G needle, employing the SYTO9 vs. PI staining method. This method has the significant advantage of providing results in less time: 15 min compared to 45 min for the DAPI method. In addition, it allows the sorting of live cells for downstream processes such as single-cell passaging approaches or mRNA analyses. Cells stained with PI (cells with damaged cell membranes and cell walls) can be excluded from the vital cells by cell sorting. However, a disadvantage of this method is its lower resolution in distinguishing cell types and the need to process live cells immediately without delay.

As an example, SYTO9 vs. PI combination differentiated six subpopulations within *B. subtilis* at sites 1 and 5, including 3 spore types, 2 vegetative cell types, as well as a group of dead cells (Figure 4). The dead cells showed only red PI relative fluorescence intensity [rel. FI] while the spores showed low to medium green SYTO9 [rel. FI] compared to the vegetative cells inhabiting the rightmost gates in the 2D plots. By back-gating two of the spore subpopulations (sp1 and sp3) with their FSC vs. SYTO9 distribution, we found further discrimination of spore types that differed in their ability to take up SYTO9 (3 with high and 2 with low SYTO9 [rel. FI]; SI Figure 7) and also differed in FSC (not shown). Similar to previous observations, a high percentage of spores were detected at sites 1–3 using the 200 µL tip (27%, 24%, and 65%, respectively), with a reduction to 10% and 15% at sites 4 and 5, respectively. In contrast, the number of vegetative cells increased from site 1 to site 5 (Figure 3), while the number of dead cells decreased (SI Figure 7). These results, along with data from the other four strains, are summarized in SI Figures 7–11.

**Figure 4:**
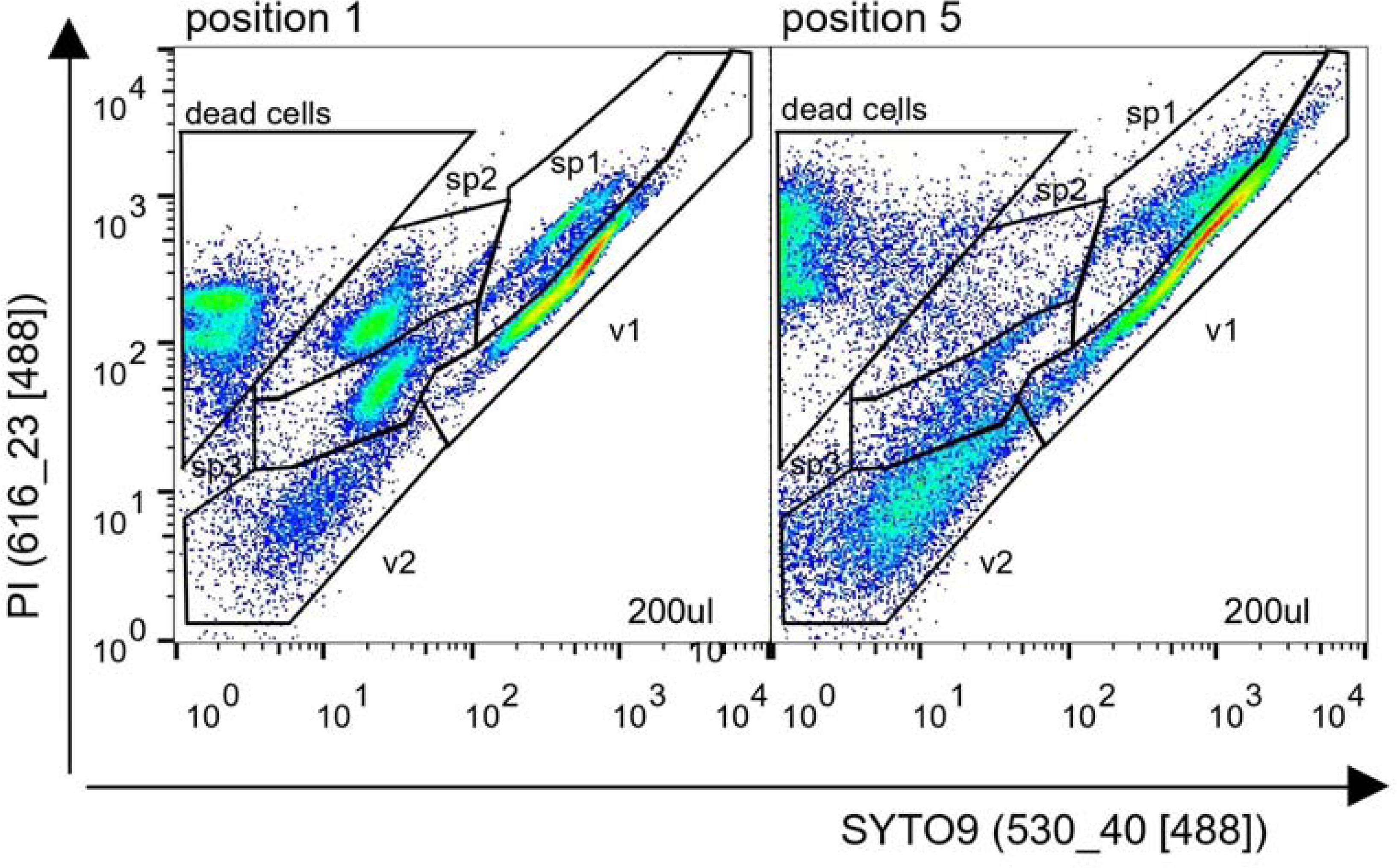
Cytometric SYTO9 vs. PI 2D plots from biopsy sites 1 and 5 of a B. subtilis colony using the 200 µL tip. The 2D SYTO9 vs. PI 2D plots reveal 6 subpopulations (2 vegetative cell types and 3 spore types, 1 dead cell type) at sites 1 and 5 but with fewer cell numbers in the spore gates at site 5.

### EFFECT OF BIOPSY SIZE PER BIOPSY SITE ON THE COMPARABILITY OF SPECIMENS

A comparative analysis of biopsy samples was conducted using the 27G needle and 10 µL to 1000 µL tips from five different sites of a colony. The samples were stained by the methods developed in this study using FSC vs. DAPI and SYTO9 vs. PI 2D plot cytometric fingerprinting. This comparison was made for each of the five different bacterial strains (Figure 5, SI Figures 13-16). Knowing that staining equilibria are highly sensitive to variations in cell/dye ratios, the objective was to detect outliers in the 2D patterns related to sample size to exclude unrepresentative biopsy data from the analysis and to find the optimal biopsy size for each strain.

**Figure 5:**
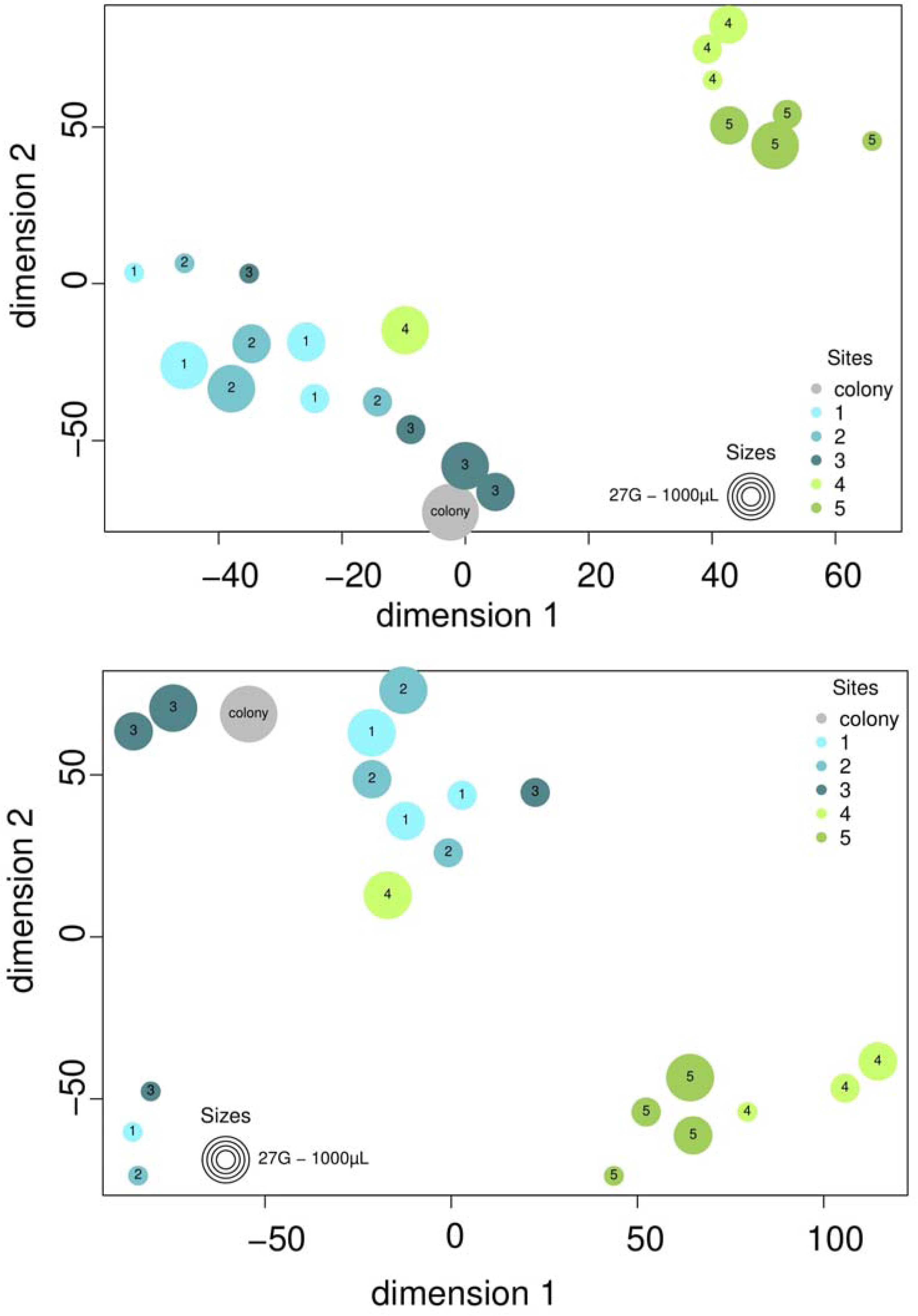
Similarity comparison of biopsy sites sampled from a 3-day-old *B. subtilis* colony using different needle or tip sizes. Samples were collected from five distinct positions within the colony (labeled 1 to 5 in respective colors, as detailed in Table 2), and measured by FCM to create cytometric fingerprints (2D plots). The size of the points reflects the dimensions of the sampling tips: 27G needle, as well as 10 µL, 200 µL, and 1000 µL pipette tips. Gray point: whole colony. Above: Samples were fixated and stained by the DAPI staining method. Below: Live samples were stained with the SYTO 9 vs. PI staining method. The cell numbers in the gates of the 2D plots were transformed by the Box-Cox transformation method and visualized by the R package t-SNE.

**Figure 6:**
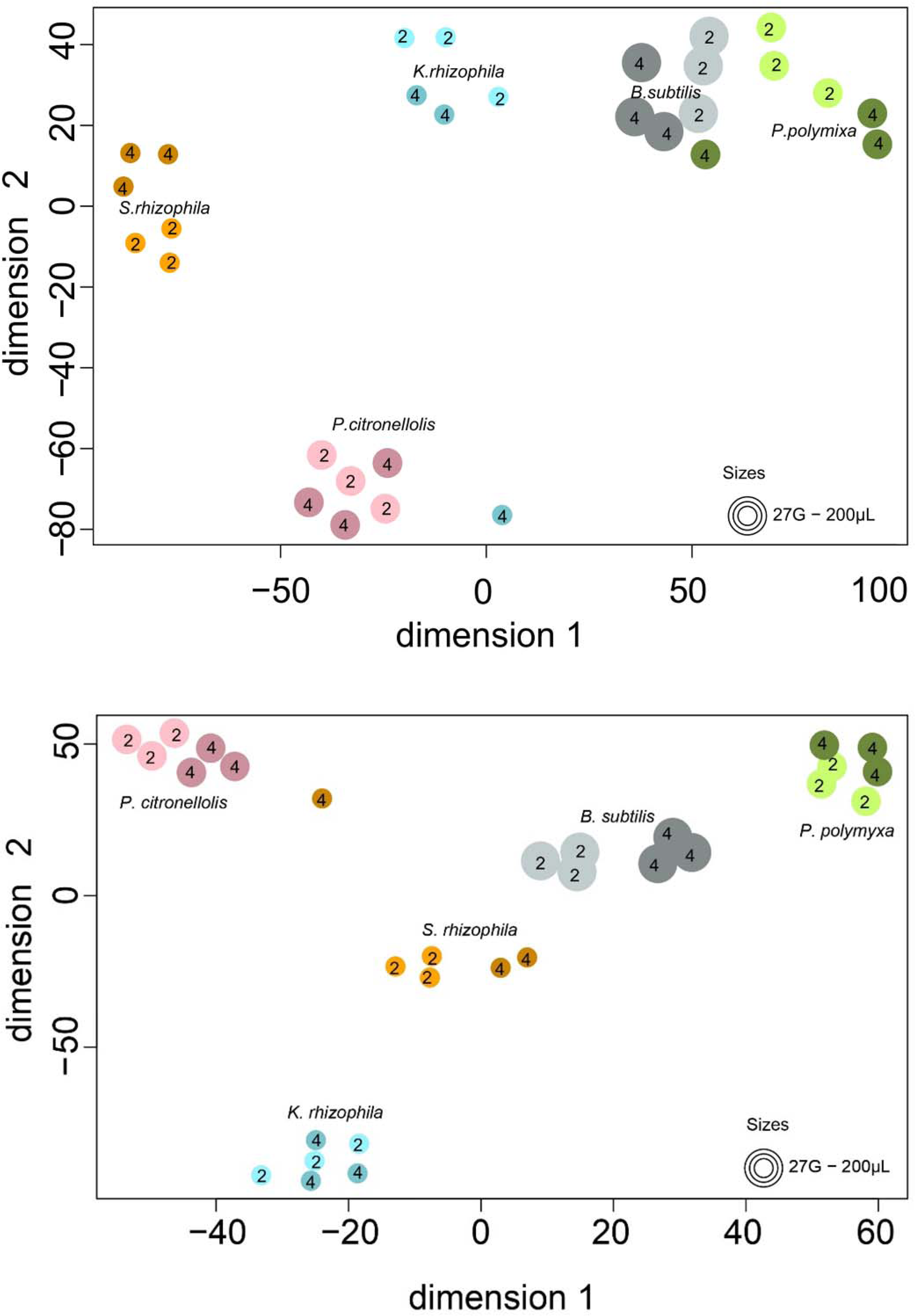
Comparison of the similarity of biopsy samples taken at the same distances from the center of respective colonies from B. subtilis, P. polymyxa, K. rhizophila, S. rhizophila, and P. citronellolis, using grid-gating. A dot represents a biopsied sample and the color of the dot represents the respective strains. The location of the sample is indicated by the shades of the color (light: site 2; dark: site 4) and the size of the needle/tips is indicated by the size of the circles. Above: FSC vs. DAPI analysis. Below: SYTO9 vs. PI analysis.

The analysis of the *B. subtilis* biopsies revealed two clear key findings: The fingerprints of the biopsies differed from the inside to the outside of the colony. The FSC vs. DAPI 2D plots (Figure 5, above) showed high clustering of the biopsy samples taken from identical locations (sites 1 - 5) in the colony when taken with different needle/tip sizes. Only one sample out of 20 (site 4, 1000 µL tip) deviated from the expected range. Similarly, the SYTO9 vs. PI 2D plots (Figure 5, below) showed a comparable clustering of local samples taken with different needle/tip sizes, except site 4 as an outlier (again, the 1000 µL tip). Interestingly, samples taken with the 27G needle showed partial clustering away from their associated locations. Therefore, second, the results suggest that the 10 µL and 200 µL tips were optimal for obtaining biopsies from this strain. The whole *B. subtilis* colony was also analyzed by FCM as a control and reflected the average distribution of the different cell types within the colony fingerprint.

Essentially, the results of the study indicate that different optimal needle/tip sizes should be used for the five strains, as larger tips are required for solid colonies, for example, to obtain enough cells for a repeatable fingerprint, while smaller tips provide enough cells from mucilaginous colonies. For *P. polymyxa*, *S. rizophila*, and *P. citronellolis* the samples taken with the different sizes of tips clustered mostly together (t-SNE plots, SI Figures 13-16). Once again, the location of the biopsy within the colony was consistently shown to be a significant factor influencing the structural characteristics of the biopsy. Specifically, for *P. polymyxa*, the use of the 27G needle is not recommended, as biopsies obtained using this needle appeared as outliers across all sites. *P. polymyxa* colonies have, similarly to *B. subtilis,* a firmer structure and yield too few cells with the 27G needle. It is recommended to sample this strain with the 10 µL tip which is also recommended for *P. citronellolis*. *S. rhizophila* and *K. rhizophila* can be biopsied with the 27G needle. Among the five studied strains, *K. rhizophila* was an exception, as less clustering was observed in its biopsy samples, suggesting that the colony structure of the strain may be more homogeneously organized under the applied conditions so that the effects of biopsy size on sample resolution cannot be followed (SI Figure 14).

### PRECISION OF THE BIOPSY PROCEDURE

As described above, the location of the biopsy within a colony is the primary determinant of the cell type composition within a sample. To verify that all biopsy samples taken from the same distance relative to the center of the colony have the same composition, triplicate samples were collected along the same concentric ring at sites 2 and 4, respectively.

The data shows that the strains have very different structures, both when analyzing FSC vs. DAPI and SYTO9 vs. PI. In addition, for most of the strains, there are also variations between the position 2 and the position 4. *K. rhizophila* shows the least variations as before and also *P. citronellolis* is less differentiated when only considering the FSC vs. DAPI analysis. But in all other tests, there was a clear separation of the population structures between sites 2 and 4, with high similarity between the triplicate samples in each instance. The results confirmed that biopsy samples obtained at equivalent distances from the colony center, displayed consistent cell type compositions, thereby validating the reliability of the biopsy procedure across the different strains and sampling tools used.

### LOCAL DIVERSIFICATION OF BACTERIAL COLONY STRUCTURES

This study shows that the local diversification of bacterial colony structures can be very high and up to 13 subpopulations with different proportions of cell types, e.g. in *B. subtilis*, can be distinguished at different locations in a colony. In contrast, there are also colony structures with a very small number of subpopulations, such as in *K. rhizophila* when stained with SYTO9 vs. PI (3 subpopulations), and their proportions hardly changed at the different biopsy sites of a colony (SI Figure 9). In *B. subtilis*, it was found that spore production was mainly localized in biopsy sites 1 to 3, while almost no spores were detected at site 5 (Figure 3). Cell cycling activity was higher in sites 4 and 5, which correlates with the high fluorescence intensity of SYTO9-stained vegetative cells observed in the same sites (Figure 7, SI Figure 17). *B. subtilis* showed six highly differentiated developmental stages of spores in both staining methods, reaching values between <1% to >15% per spore type. In *P. polymyxa,* spore production predominantly occurred at position 2, though at a significantly lower overall abundance compared to other positions. Notably, only spore type sp4 was present at a relatively higher abundance, reaching up to 6%. Furthermore, the cell cycle activity was most pronounced at position 5 (Figure 7), where over 50% of the cells exhibited more than double the chromosomal content, as indicated by c2xn DAPI staining (SI Figure 18). The other three strains, *K. rhizophila*, *S. rhizophila*, and *P. citronellolis* did not show any spores as they are no spore producers. In *K. rhizophila*, the cell cycle was performed at all sites (Figure 7), but mainly by the c1n, c2n, and c4n cells (DAPI-stained, SI Figure 19), which was also reflected in the high proportions of vegetative SYTO9-stained cells (Figure 7). Growth was relatively evenly distributed across all regions of the colony. A similar pattern was observed in *S. rhizophila* (Figure 7); however, upon closer examination (SI Figure 20), the highest growth activity was concentrated at position 5 (c2n-c). In contrast, the highest proportion of cells containing a single chromosome was identified in positions 1 and 2. (c1n-b; SI Figure 20). For the SYTO9-stained cells, the highest proportions of vegetative cells were observed at positions 4 and 5 (v1). In *P. citronellolis,* the abundance of actively growing cells was relatively low, with less than 25% detected at each position (Figure 7). These growing cells were evenly distributed across the five locations, though higher proportions were observed in the c2n cell population (SI Figure 21). Unlike other strains, *P. citronellolis* efficiently expelled the fluorescent dyes from its cells within a few minutes, resulting in a notably high percentage of less-stained cells, exceeding 75% in DAPI staining (Figure 7). Furthermore, over 80% of the SYTO9-stained cells exhibited low fluorescence intensity (ls cells) and were therefore not included in the SYTO9 boxplots presented in Figure 7.

**Figure 7.**
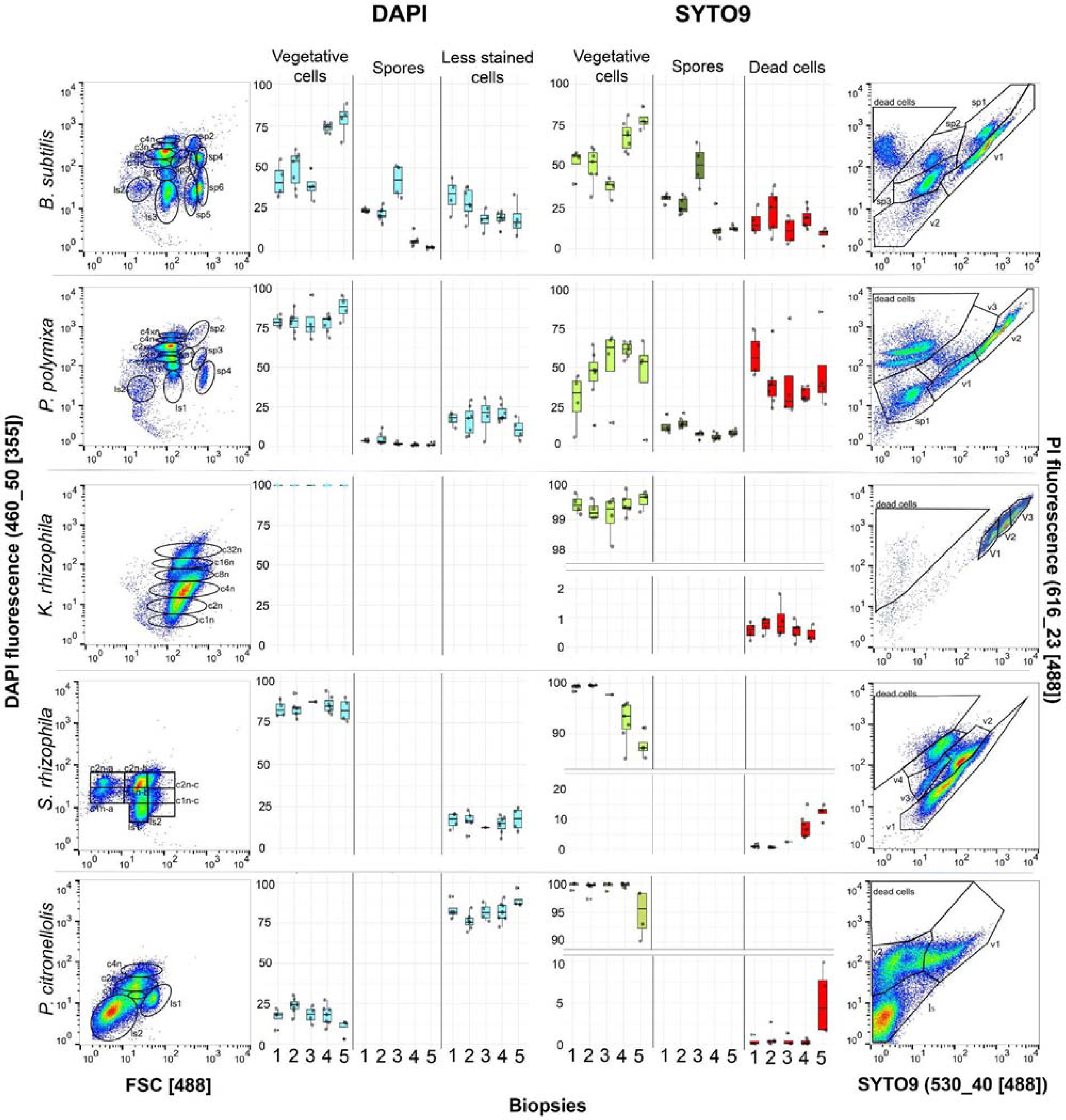
Proportions of cell types of *B. subtilis, P. polymyxa, K. rhizophila, S. rhizophila*, and *P. citronellolis* at the five biopsy sites of colonies. Left: FSC vs. DAPI analysis. Right: SYTO9 vs. PI analysis. All 2D plots show different cell types which were gated according to physiological characteristics into subpopulations for relative cell abundances assessment. Specifically, vegetative cells (cxn), spores (sp) and less stained cells (ls) for DAPI staining (all in blue); and vegetative cells (v; light green), spores (sp; dark green) and dead cells (red) for SYTO9 vs. PI staining were discriminated.

The proportion of dead cells was high in *B. subtilis* (over 10%) and *P. polymyxa* (exceeding 25%), while for the other three strains, it remained around 1%. However, at site 5, the proportion of dead cells increased to approximately 10% for *S. rhizophila* and *P. citronellolis*, suggesting that actively growing cells are either more susceptible to PI staining or more readily take up the PI dye, consistent with previous observations in other strains (Shi et al., 2007). The least variability in cytometric fingerprint patterns was observed for *K. rhizophila*. Furthermore, both cytometric methods demonstrated excellent discriminatory capabilities across all strains, providing identified cell types within 45 minutes (FSC vs. DAPI) and 15 minutes (SYTO9 vs. PI), respectively.

## DISCUSSION

On undisturbed surfaces where nutrient, carbon, and energy sources are evenly distributed, a single bacterial cell typically grows into a circular-shaped colony. The diameter, height, and macro-structure of the colony depend on both the type of strain and the growth conditions (Chacón et al., 2018; Xue et al., 2021). Additionally, the ability to produce coloration and extracellular polymeric substances (EPS) also varies between strains (SI Figure 1) and is often used as a quick way to assess the purity of strains cultured on agar plates (Amoretti et al., 2002). Many strains show more or less textured patterns of varying concentric circles, as well as valleys and mounds (Ben-Jacob et al., 2000). Although all cells growing in a colony are genetically identical and originated from the same ancestor, physiological heterogeneity typically emerges after a few generations, often observable within three days. Also, the mechanical properties of bacteria and surface have an important influence on colony structure (Araújo et al., 2019; Zheng et al., 2021), such as the formation of channels that facilitate nutrient distribution among different cell types.

Despite being uniform clusters of genetically identical cells, these variable multicellular structures reveal interactions between individual cells and their abiotic environment. As early as 1988, James Shapiro highlighted the multicellular nature of bacterial colonies, proposing that individual bacteria do not act autonomously but as a collective (Shapiro, 1988). Properties of bacteria are usually investigated on a bulk scale, assuming homogeneity in cellular behavior, which is the standard for biotechnological processes. However, it has long been recognized that even in planktonic cultures, physiological heterogeneity exists between cell types, driven by factors such as cell cycle states, age distributions, mutations, or transcriptional noise (Ackermann, 2015; Bergmiller & Ackermann, 2011; Müller et al., 2010; Süel et al., 2007). James Shapiro suggested that other parameters in a colony are also important for functional diversity among cells: location and time. He rejected the inheritance of cell types as the cause of the differentiation within colonies and instead suggested that interaction mechanisms were caused by temporal control of development (Shapiro, 1988). A very prominent example of such a development is the EPS production of some cells of a colony to promote matrix production for adherence to a surface while others act as benefitting cheaters. Demonstrating that isogenic populations can diversify into phenotypically distinct variants (Dragoš et al., 2018), examples among bacteria have shown that cooperative matrix producers responded by increasing gene expression for matrix production, thereby limiting the number of cheaters (Amoretti et al., 2002; Martin et al., 2020; Nadell et al., 2016; Nadell & Bassler, 2011). The number and geographical distribution of isogenic founder cells were also found to influence the morphology and behavior of the mature colony (Eigentler et al., 2022). The authors call it a race-for-space, where competition for nutrients dominates the dynamics of cell growth and the resulting spatial structure and relative densities in the mature biofilm (Eigentler et al., 2022).

Bacterial colonies are ecologically complex and heterogeneous populations in which biogeography drives cellular differentiation and specialization. The spatial heterogeneity observed within colonies, such as the differential cell cycle activities and spore formation across different colony regions, suggests that bacterial cells modulate their physiological states according to their local microenvironment (Huang et al., 2024). *B. subtilis* is a model organism for studying the transition from planktonic cells to physiological highly differentiated cells in a colony (Arnaouteli et al., 2021). This strain is therefore also a versatile and beneficial producer strain that is widely used in biotechnological applications, including the production of stabilizing multiphase formulations, calcite precipitation, food fermentation, or as a probiotic (Arnaouteli et al., 2021; Bromley & MacPhee, 2017; Marzorati et al., 2020; Reeksting et al., 2020). The present study confirms the high degree of physiological heterogeneity of the strain and also diversification as a function of its spatial location in a colony. The generally high number of dead cells in the inner circle of a colony (>23%) and the accumulation of spores especially at site 3 of a *B. subtilis* colony (>51%) are particularly noteworthy. In contrast, increased growth behavior was observed at the outer edge of the colonies due to the increased number of chromosomes (>74%). The observed cell cycle activity with higher proliferation at the edges of the colony may reflect a strategy to exploit the abundant nutrients and oxygen at the edges. In contrast, cells at the center of a colony adapt their behavior and physiology to the limited local availability of resources, following ecological principles of resource allocation and niche differentiation. Localized sporulation in *B. subtilis*, particularly in the inner regions of the colony, is an evolutionary strategy to protect the population from nutrient deprivation or other adverse conditions (Lopez et al., 2008).

The other four strains used in the study have been studied much less or not at all in terms of their heterogeneity and biogeographical behavior within a colony. All strains have fully sequenced genomes. *K. rhizophila* and S*. rhizophila* were isolated from natural sources, such as the rhizosphere of the soil (Alavi et al., 2014; Takarada et al., 2008), and were used to test for antibiotics in food and medical samples (Dafale et al., 2013, Elhosieny et al., 2023). In contrast to *B. subtilis*, the gram-positive *K. rhizophila,* and the gram-negative *S. rhizophila* showed growth activity at virtually all sites of the colony, with at least *S. rhizophila* showing a slight increase in activity at site 5, analogous to *B. subtilis*. The number of dead cells was generally low, although *S. rhizophila* showed an increase in cell death at site 5, where no cell death was anticipated due to favorable growth conditions. *P. polymyxa* is a spore producer and motile strain of soil origin (He et al., 2007). The strain exhibited unexpectedly high numbers of PI-stained cells (>45%) in the center of the colony, accompanied by low spore formation. Whether these PI-stained cells were really dead or whether they had taken up PI due to open membranes and cell walls during growth, as was also observed for other strains (Shi et al., 2007), was not investigated in this study. A different behavior was observed for *P. citronellosis*. This strain is a motile gamma-proteobacterium, was also isolated from the rhizosphere (Remus-Emsermann et al., 2016) and contains the chromosomally located red fluorescent protein mScarlet-I (Schlechter et al., 2018) and shows a permanent red fluorescence after excitation with the 561 nm laser (Bindels et al., 2017), but also to a lesser extent with the 488 nm laser (not shown). Due to the overlap of the mScarlet-I emission with the PI emission, it was not possible to distinguish between live and dead cells. In addition, this strain exhibited high fluorescent dye pumping activities, requiring strict time-dependent flow cytometric measurements. Growth activities were evenly distributed between sites of the colony.

Therefore, some bacterial strains, such as *K. rhizophila* and *P. citronellolis* showed uniform growth activities across different regions of the colony, while others showed higher activities at specific locations, e.g. at site 5. The uniform behavior was particularly pronounced for the *K. rhizophila* strain. The assumption that it is not the method that is unspecific, but rather that this strain imparts little structure to its colony, is supported by the t-distributed Stochastic Neighbor Embedding plot (t-SNE plots, SI Figure 14), where different biopsy sizes taken from different colony positions showed a rather random distribution when treated with both staining methods. In contrast, the t-SNE plots for the strains *B. subtilis*, *P. polymyxa, S. rhizophila,* and even *P. citronellolis* demonstrated clear clustering of samples from all other sites for the selected tips or needle, emphasizing that the developed methods are suitable and effective for analyzing the biogeography of bacterial colony. The developed methods enable rapid determination of sample heterogeneity within 45 min (FSC vs. DAPI) and 15 min (SYTO9 vs. PI) and can be universally applied.

Expanding these methods to previously unstudied strains is essential to unravel the richness of bacterial lifestyles that may have developed different and previously unknown strategies to manage the complex process of competition and cooperation within a biofilm colony. Among the five strains, *B. subtilis* has already been studied extensively because of its compartmentalization and the different functions of cell types at specific positions in a colony (Ackermann, 2015; Dragoš et al., 2018; Martin et al., 2020). Recently, a new concept has been published that favors a clock and wavefront mechanism to explain ring-shaped structures in colonies of *B. subtilis* (Chou et al., 2022). This concept is not only based on the development of heterogeneous cell states due to different nutrient availability or noise in gene regulation (Arnaouteli et al., 2021; Chai et al., 2008; Maamar et al., 2007; Setlow & Christie, 2023), but also on cellular differentiation and organization caused by cell-autonomous oscillations. The authors used nitrogen limitation to generate a deterministic clock and wavefront mechanism that appears to allow local amplification of the nitrogen stress response, resulting in spore formation not only within the colony but also near the colony edges, despite nitrogen was still present. This led to segmented spore formation in the colony (Chou et al., 2022), which was also observed in *B. subtilis* and *P. polymyxa* colonies at sites 3 and 2, respectively, in this study. This behavior is analogous to multicellular organisms such as for vertebrate somitogenesis where clock and wavefront mechanisms cause spatially recurring segments (Hubaud et al., 2017). Collective oscillatory behavior has also been reported for other microorganisms. For example, *Neisseria gonorrhoeae* exhibits collective polarization dynamics in response to local oxygen concentration in older colonies (Hennes et al., 2023). In contrast, this study found minimal structural diversification and low cell type heterogeneity in *K. rhizophila* colonies. Vassallo et al. (2019) proposed a model in which nutrient concentration and diffusion were shown to have a major influence on colony morphology and structure. This extends to the potential functions of individual cell types within a colony and the overall functionality of a colony as an integrated system. Therefore, a lower degree of cell heterogeneity can be expected if a colony itself is less structured (Claessen et al., 2014; Nadell et al., 2010; Puri et al., 2023).

## CONCLUSION

The study indicates how cell populations specialize within different regions of a colony, to guarantee efficient growth and survival at the micro-scale. Taking biopsies of isogenic bacterial colonies at defined sites and analyzing the phenotypic heterogeneity of these cells using FCM could serve as a valuable tool to decipher the ecological properties of cell types in response to growth conditions such as carbon and energy resources, nutrients, or biophysical parameters. The method can also be used for the study of local cell responses to stress, as can be the response to colony injury (Ye et al., 2024) or toxic components such as antibiotics (Ma et al., 2023). In the future, cell sorting of individual cells from colony biofilms, e.g. sc-mRNA analysis will provide even deeper insight into the regulatory and metabolic processes within microbial populations. Understanding the local diversification of colonies and the respective local cell functions can also help to understand the overarching function of whole colonies.

## DATA AVAILABILITY STATEMENT

The raw flow cytometry data is available in the open repository http://zenodo.org and the data can be accessed by the reviewers via the following link: https://doi.org/10.5281/zenodo.13791365

All other data used in the manuscript are available as Supplementary Information SI Figures 1-21.

## Acknowledgments

This work was supported by the Deutsche Forschungsgemeinschaft (DFG) within the SPP 2389 priority program “Emergent Functions of Bacterial Multicellularity” (Grant number: 503905203)

## Author contribution statement

G.A., K.S., and S.M. conceived the ideas and designed the methodology; K.S. and G.A. collected the data; G.A. and K.S. analyzed the data; S.M. led the writing of the manuscript. All authors contributed critically to the drafts and gave final approval for publication.

## REFERENCES

Ackermann, M. (2015). A functional perspective on phenotypic heterogeneity in microorganisms. Nature Reviews Microbiology, 13(8), 497–508. 10.1038/nrmicro3491

Alavi, P., Starcher, M. R., Thallinger, G. G., Zachow, C., Müller, H., & Berg, G. (2014). *Stenotrophomonas* comparative genomics reveals genes and functions that differentiate beneficial and pathogenic bacteria. BMC Genomics, 15(1), 1–15. 10.1186/1471-2164-15-482

Amoretti, M., Amsler, C., Bonomi, G., Bouchta, A., Bowe, P., Carraro, C., Cesar, C. L., Chaliton, M., Collier, M. J. T., Doser, M., Filippini, V., Fine, F. S., Fontana, A., Fujiwara, M. C., Funakoshi, R., Genova, P., Hangst, J. S., Hayano, R. S., Holzscheiter, M. H., … Van Der Werf, D. P. (2002). Production and detection of cold antihydrogen atoms. Nature, 419(6906), 456–459. 10.1038/nature01096

Araújo, G. R. d. S., Viana, N. B., Gómez, F., Pontes, B., & Frases, S. (2019). The mechanical properties of microbial surfaces and biofilms. Cell Surface, 5, 100028. 10.1016/j.tcsw.2019.100028

Arnaouteli, S., Bamford, N. C., Stanley-Wall, N. R., & Kovács, Á. T. (2021). *Bacillus subtilis* biofilm formation and social interactions. Nature Reviews Microbiology, 19(9), 600–614. 10.1038/s41579-021-00540-9

Atkinson, A. C., Riani, M., & Corbellini, A. (2021). The Box–Cox Transformation: Review and Extensions. Statistical Science, 36(2), 239–255. 10.1214/20-STS778

Bar-On, Y. M., Phillips, R., & Milo, R. (2018). The biomass distribution on Earth. Proceedings of the National Academy of Sciences of the United States of America, 115(25), 6506–6511. 10.1073/pnas.1711842115

Bellais, S., Nehlich, M., Ania, M., Duquenoy, A., Mazier, W., van den Engh, G., Baijer, J., Treichel, N. S., Clavel, T., Belotserkovsky, I., & Thomas, V. (2022). Species-targeted sorting and cultivation of commensal bacteria from the gut microbiome using flow cytometry under anaerobic conditions. Microbiome, 10(1), 1–17. 10.1186/s40168-021-01206-7

Ben-Jacob, E., Cohen, I., & Levine, H. (2000). Cooperative self-organization of microorganisms. Advances in Physics, 49(4), 395–554. 10.1080/000187300405228

Bergmiller, T., & Ackermann, M. (2011). Pole age affects cell size and the timing of cell division in *Methylobacterium extorquens* AM1. Journal of Bacteriology, 193(19), 5216–5221. 10.1128/JB.00329-11

Bien, T., Koerfer, K., Schwenzfeier, J., Dreisewerd, K., & Soltwisch, J. (2022). Mass spectrometry imaging to explore molecular heterogeneity in cell culture. Proceedings of the National Academy of Sciences of the United States of America, 119(29), 1–12. 10.1073/pnas.2114365119

Bindels, D. S., Haarbosch, L., van Weeren, L., Postma, M., Wiese, K. E., Mastop, M., Aumonier, S., Gotthard, G., Royant, A., Hink, M. A., & Gadella, T. W. J. J. (2017). mScarlet: a bright monomeric red fluorescent protein for cellular imaging. Nature Methods, 14(1), 53–56. 10.1038/nmeth.4074

Bingham, E. P., & Ratcliff, W. C. (2024). A nonadaptive explanation for macroevolutionary patterns in the evolution of complex multicellularity. Proceedings of the National Academy of Sciences, 121(7), e2319840121. 10.1073/pnas.2319840121

Box, G. E. P., & Cox, D. R. (1964). An analysis of transformations. Journal of the Royal Statistical Society. Series B (Methodological), 26, 211–243, 10.1111/j.2517-6161.1964.tb00553.x</boxcap>

Branda, S. S., González-Pastor, J. E., Ben-Yehuda, S., Losick, R., & Kolter, R. (2001). Fruiting body formation by *Bacillus subtilis*. Proceedings of the National Academy of Sciences, 98(20), 11621– 11626. 10.1073/pnas.191384198

Brockmann, E. U., Steil, D., Bauwens, A., Soltwisch, J., & Dreisewerd, K. (2019). Advanced methods for MALDI-MS imaging of the chemical communication in microbial communities. Analytical Chemistry, 91(23), 15081–15089. 10.1021/acs.analchem.9b03772

Bromley, K. M., & MacPhee, C. E. (2017). BslA-stabilized emulsion droplets with designed microstructure. Interface Focus, 7(4), 20160124. 10.1098/rsfs.2016.0124

Chacón, J. M., Möbius, W., & Harcombe, W. R. (2018). The spatial and metabolic basis of colony size variation. ISME Journal, 12(3), 669–680. 10.1038/s41396-017-0038-0

Chai, Y., Chu, F., Kolter, R., & Losick, R. (2008). Bistability and biofilm formation in *Bacillus subtilis*. Molecular Microbiology, 67(2), 254–263. 10.1111/j.1365-2958.2007.06040.x

Chou, K.-T., Lee, D. D., Chiou, J., Galera-Laporta, L., Ly, S., Garcia-Ojalvo, J., & Süel, G. M. (2022). A segmentation clock patterns cellular differentiation in a bacterial biofilm. Cell, 185(1), 145–157.e13. 10.1016/j.cell.2021.12.001

Cichocki, N., Hübschmann, T., Schattenberg, F., Kerckhof, F. M., Overmann, J., & Müller, S. (2020). Bacterial mock communities as standards for reproducible cytometric microbiome analysis. Nature Protocols, 15(9), 2788–2812. 10.1038/s41596-020-0362-0

Claessen, D., Rozen, D. E., Kuipers, O. P., Søgaard-Andersen, L., & Van Wezel, G. P. (2014). Bacterial solutions to multicellularity: A tale of biofilms, filaments and fruiting bodies. Nature Reviews Microbiology, 12(2), 115–124. 10.1038/nrmicro3178

Dafale, N. A., Agarwal, P., Semwal, U. P., & Singh, G. N. (2013). Development and validation of microbial bioassay for the quantification of potency of the antibiotic cefuroxime axetil. Analytical Methods, 5, 690–698. 10.1039/C2AY25848J

Dalziel, B. D., Novak, M., Watson, J. R., & Ellner, S. P. (2021). Collective behaviour can stabilize ecosystems. Nature Ecology and Evolution, 5(10), 1435–1440. 10.1038/s41559-021-01517-w

Dragoš, A., Lakshmanan, N., Martin, M., Horváth, B., Maróti, G., García, C. F., Lieleg, O., & Kovács, Á. T. (2018). Evolution of exploitative interactions during diversification in *Bacillus subtilis* biofilms. FEMS Microbiology Ecology, 94(1), 1–12. 10.1093/femsec/fix155

Drescher, K., Dunkel, J., Nadell, C. D., Van Teeffelen, S., Grnja, I., Wingreen, N. S., Stone, H. A., & Bassler, B. L. (2016). Architectural transitions in *Vibrio cholerae* biofilms at single-cell resolution. Proceedings of the National Academy of Sciences of the United States of America, 113(14), E2066– E2072. 10.1073/pnas.1601702113

Eigentler, L., Kalamara, M., Ball, G., MacPhee, C. E., Stanley-Wall, N. R., & Davidson, F. A. (2022). Founder cell configuration drives competitive outcome within colony biofilms. ISME Journal, 16(6), 1512–1522. 10.1038/s41396-022-01198-8

Elhosieny, A. A. E., Zayed, M. S., Selim, S. M., Yassen, A. M., & Abdel Aziz, N. H. (2023). *Stenotrophomonas rhizophila* a novel plant-associated bacterium with distinguished PGPRs properties. Arab Universities Journal of Agricultural Sciences, 31(1), 41–50. 10.21608/ajs.2023.159562.1493

Fisher, R. M., Cornwallis, C. K., & West, S. A. (2013). Group formation, relatedness, and the evolution of multicellularity. Current Biology, 23(12), 1120–1125. 10.1016/j.cub.2013.05.004

Flemming, H. C., & Wuertz, S. (2019). Bacteria and archaea on Earth and their abundance in biofilms. Nature Reviews Microbiology, 17(4), 247–260. 10.1038/s41579-019-0158-9

Freese, E., Heinze, J. E., & Galliers, E. M. (1979). Partial purine deprivation causes sporulation of *Bacillus subtilis* in the presence of excess ammonia, glucose and phosphate. Microbiology, 115(1), 193–205. 10.1099/00221287-115-1-193

Freiherr von Boeselager, R., Pfeifer, E., & Frunzke, J. (2018). Cytometry meets next-generation sequencing – RNA-Seq of sorted subpopulations reveals regional replication and iron-triggered prophage induction in *Corynebacterium glutamicum*. Scientific Reports, 8(1), 1–13. 10.1038/s41598-018-32997-9

Haange, S. B., Jehmlich, N., Krügel, U., Hintschich, C., Wehrmann, D., Hankir, M., Seyfried, F., Froment, J., Hübschmann, T., Müller, S., Wissenbach, D. K., Kang, K., Buettner, C., Panagiotou, G., Noll, M., Rolle-Kampczyk, U., Fenske, W., & Von Bergen, M. (2020). Gastric bypass surgery in a rat model alters the community structure and functional composition of the intestinal microbiota independently of weight loss. Microbiome, 8(1), 1–17. 10.1186/s40168-020-0788-1

Hartmann, R., Jeckel, H., Jelli, E., Singh, P. K., Vaidya, S., Bayer, M., Rode, D. K. H., Vidakovic, L., Díaz-Pascual, F., Fong, J. C. N., Dragoš, A., Lamprecht, O., Thöming, J. G., Netter, N., Häussler, S., Nadell, C. D., Sourjik, V., Kovács, Á. T., Yildiz, F. H., & Drescher, K. (2021). Quantitative image analysis of microbial communities with BiofilmQ. Nature Microbiology, 6(2), 151–156. 10.1038/s41564-020-00817-4

Hartmann, R., Singh, P. K., Pearce, P., Mok, R., Song, B., Díaz-Pascual, F., Dunkel, J., & Drescher, K. (2019). Emergence of three-dimensional order and structure in growing biofilms. Nature Physics, 15(3), 251–256. 10.1038/s41567-018-0356-9

Hennes, M., Bender, N., Cronenberg, T., Welker, A., & Maier, B. (2023). Collective polarization dynamics in bacterial colonies signify the occurrence of distinct subpopulations. PLoS Biology, 21(1), 1–20. 10.1371/journal.pbio.3001960

Hinton, G. E., & Roweis, S. T. (2002). Stochastic Neighbor Embedding. Neural Information Processing Systems. https://api.semanticscholar.org/CorpusID:20240

Höfler, C., Heckmann, J., Fritsch, A., Popp, P., Gebhard, S., Fritz, G., & Mascher, T. (2016). Cannibalism stress response in *Bacillus subtilis*. Microbiology (United Kingdom), 162(1), 164–176. 10.1099/mic.0.000176

Huang, S.-W., Lim, S.-K., Yu, Y.-A., Pan, Y.-C., Lien, W.-J., Mou, C.-Y., Hu, C.-M. J., & Mou, K. Y. (2024). Overcoming the nutritional immunity by engineering iron-scavenging bacteria for cancer therapy eLife 12:RP90798. 10.7554/eLife.90798.3

Hubaud, A., Regev, I., Mahadevan, L., & Pourquié, O. (2017). Excitable dynamics and Yap-dependent mechanical cues drive the segmentation clock. Cell, 171(3), 668–682.e11. 10.1016/j.cell.2017.08.043

Imdahl, F., Vafadarnejad, E., Homberger, C., Saliba, A. E., & Vogel, J. (2020). Single-cell RNA-sequencing reports growth-condition-specific global transcriptomes of individual bacteria. Nature Microbiology, 5(10), 1202–1206. 10.1038/s41564-020-0774-1

Jehmlich, N., Hübschmann, T., Gesell Salazar, M., Völker, U., Benndorf, D., Müller, S., Von Bergen, M., & Schmidt, F. (2010). Advanced tool for characterization of microbial cultures by combining cytomics and proteomics. Applied Microbiology and Biotechnology, 88(2), 575–584. 10.1007/s00253-010-2753-6

Khan, M. S. I., Oh, S. W., & Kim, Y. J. (2020). Power of scanning electron microscopy and energy dispersive X-Ray analysis in rapid microbial detection and identification at the single cell level. Scientific Reports, 10(1), 1–10. 10.1038/s41598-020-59448-8

Koch, C., Günther, S., Desta, A. F., Hübschmann, T., & Müller, S. (2013). Cytometric fingerprinting for analyzing microbial intracommunity structure variation and identifying subcommunity function. Nature Protocols, 8(1), 190–202. 10.1038/nprot.2012.149

Krijthe, J.H. (2015). Rtsne: T-Distributed Stochastic Neighbor Embedding using Barnes-Hut implementation. R package version 0.17. https://github.com/jkrijthe/Rtsne.

Liu, J., Prindle, A., Humphries, J., Gabalda-Sagarra, M., Asally, M., Lee, D. D., Ly, S., Garcia-Ojalvo, J., & Süel, G. M. (2015). Metabolic co-dependence gives rise to collective oscillations within biofilms. Nature, 523(7562), 550–554. 10.1038/nature14660

Longo, S. K., Guo, M. G., Ji, A. L., & Khavari, P. A. (2021). Integrating single-cell and spatial transcriptomics to elucidate intercellular tissue dynamics. Nature Reviews. Genetics, 22(10), 627– 644. 10.1038/s41576-021-00370-8

López-Gálvez, J., Schiessl, K., Besmer, M. D., Bruckmann, C., Harms, H., & Müller, S. (2023). Development of an automated online flow cytometry method to quantify cell density and fingerprint bacterial communities. Cells, 12(12). 10.3390/cells12121559

Lopez, D., Vlamakis, H., & Kolter, R. (2008). Generation of multiple cell types in *Bacillus subtilis*. FEMS Microbiology Reviews, 33(1), 152–163. 10.1111/j.1574-6976.2008.00148.x

Lyons, N. A., & Kolter, R. (2015). On the evolution of bacterial multicellularity. Current Opinion in Microbiology, 24, 21–28. 10.1016/j.mib.2014.12.007

Ma, P., Amemiya, H. M., He, L. L., Gandhi, S. J., Nicol, R., Bhattacharyya, R. P., Smillie, C. S., & Hung, D. T. (2023). Bacterial droplet-based single-cell RNA-seq reveals antibiotic-associated heterogeneous cellular states. Cell, 186(4), 877–891.e14. 10.1016/j.cell.2023.01.002

Maamar, H., Raj, A., & Dubnau, D. (2007). Noise in gene expression determines cell fate in *Bacillus subtilis*. Science (New York, N.Y.), 317(5837), 526–529. 10.1126/science.1140818

Magnabosco, C., Lin, L. H., Dong, H., Bomberg, M., Ghiorse, W., Stan-Lotter, H., Pedersen, K., Kieft, T. L., van Heerden, E., & Onstott, T. C. (2018). The biomass and biodiversity of the continental subsurface. Nature Geoscience, 11(10), 707–717. 10.1038/s41561-018-0221-6

Martin, M., Dragoš, A., Otto, S. B., Schäfer, D., Brix, S., Maróti, G., & Kovács, Á. T. (2020). Cheaters shape the evolution of phenotypic heterogeneity in *Bacillus subtilis* biofilms. ISME Journal, 14(9), 2302– 2312. 10.1038/s41396-020-0685-4

Marzorati, M., Van den Abbeele, P., Bubeck, S. S., Bayne, T., Krishnan, K., Young, A., Mehta, D., & Desouza, A. (2020). *Bacillus subtilis* HU58 and *Bacillus coagulans* SC208 probiotics reduced the effects of antibiotic-induced gut microbiome dysbiosis in an M-SHIME® model. Microorganisms, 8(7), 1–15. 10.3390/microorganisms8071028

McNulty, R., Sritharan, D., Pahng, S. H., Meisch, J. P., Liu, S., Brennan, M. A., Saxer, G., Hormoz, S., & Rosenthal, A. Z. (2023). Probe-based bacterial single-cell RNA sequencing predicts toxin regulation. Nature Microbiology, 8(5), 934–945. 10.1038/s41564-023-01348-4

Milo, R., & Phillips, R. (2015). Cell Biology by the Numbers (1st ed.). Milo, R., & Phillips, R. Garland Science. 10.1201/9780429258770

Müller, S. (2007). Modes of cytometric bacterial DNA pattern: A tool for pursuing growth. Cell Proliferation, 40(5), 621–639. 10.1111/j.1365-2184.2007.00465.x

Müller, S., Harms, H., & Bley, T. (2010). Origin and analysis of microbial population heterogeneity in bioprocesses. Current Opinion in Biotechnology, 21(1), 100–113. 10.1016/j.copbio.2010.01.002

Nadell, C. D., Drescher, K., & Foster, K. R. (2016). Spatial structure, cooperation and competition in biofilms. Nature Reviews Microbiology, 14(9), 589–600. 10.1038/nrmicro.2016.84

Nadell, C. D., & Bassler, B.L. (2011). A fitness trade-off between local competition and dispersal in *Vibrio cholerae* biofilms. Proc. Natl. Acad. Sci. USA. 23; 108(34):14181-5. doi: 10.1073/pnas.1111147108.

Nadell, C. D., Foster, K. R., & Xavier, J. B. (2010). Emergence of spatial structure in cell groups and the evolution of cooperation. PLoS Computational Biology, 6(3). 10.1371/journal.pcbi.1000716

Oksanen, J., Simpson, G., Blanchet, F., Kindt, R., Legendre, P., Minchin, P., O’Hara, R., Solymos, P., Stevens, M., Szoecs, E., Wagner, H., Barbour, M., Bedward, M., Bolker, B., Borcard, D., Carvalho, G., Chirico, M., De Caceres, M., Durand, S., … Weedon, J. (2024). vegan: Community Ecology Package. R package version 2.7-0,. https://github.com/vegandevs/vegan, https://vegandevs.github.io/vegan/.

Paula, A. J., Hwang, G., & Koo, H. (2020). Dynamics of bacterial population growth in biofilms resemble spatial and structural aspects of urbanization. Nature Communications, 11(1), 1–14. 10.1038/s41467-020-15165-4

Puri, D., Fang, X., & Allison, K. R. (2023). Evidence of a possible multicellular life cycle in *Escherichia coli*. IScience, 26(1), 105795. 10.1016/j.isci.2022.105795

Reeksting, B. J., Hoffmann, T. D., Tan, L., Paine, K., & Gebhard, S. (2020). In-depth profiling of calcite precipitation by environmental bacteria reveals fundamental mechanistic differences with relevance to application. Applied and Environmental Microbiology, 86(7). 10.1128/AEM.02739-19

Remus-Emsermann, M. N. P., Schmid, M., Gekenidis, M.-T., Pelludat, C., Frey, J. E., Ahrens, C. H., & Drissner, D. (2016). Complete genome sequence of *Pseudomonas citronellolis* P3B5, a candidate for microbial phyllo-remediation of hydrocarbon-contaminated sites. Standards in Genomic Sciences, 11, 75. 10.1186/s40793-016-0190-6

Schlafer, S., & Meyer, R. L. (2017). Confocal microscopy imaging of the biofilm matrix. Journal of Microbiological Methods, 138, 50–59. 10.1016/j.mimet.2016.03.002

Schlechter, R. O., Jun, H., Bernach, M., Oso, S., Boyd, E., Muñoz-Lintz, D. A., Dobson, R. C. J., Remus, D. M., & Remus-Emsermann, M. N. P. (2018). Chromatic bacteria – A broad host-range plasmid and chromosomal insertion toolbox for fluorescent protein expression in bacteria. Frontiers in Microbiology, 9, 1–14. 10.3389/fmicb.2018.03052

Setlow, P., & Christie, G. (2023). New thoughts on an old topic: secrets of bacterial spore resistance slowly being revealed. Microbiology and Molecular Biology Reviews, 87(2). 10.1128/mmbr.00080-22

Shapiro, J. A. (1998). Thinking about bacterial populations as multicellular organisms. Annual Review of Microbiology, 52, 81–104. 10.1146/annurev.micro.52.1.81

Shi, L., Günther, S., Hübschmann, T., Wick, L. Y., Harms, H., & Müller, S. (2007). Limits of propidium iodide as a cell viability indicator for environmental bacteria. Cytometry Part A, 71(8), 592–598. 10.1002/cyto.a.20402

Süel, G. M., Kulkarni, R. P., Dworkin, J., Garcia-Ojalvo, J., & Elowitz, M. B. (2007). Tunability and noise dependence in differentiation dynamics. Science (New York, N.Y.), 315(5819), 1716–1719. 10.1126/science.1137455

Takarada, H., Sekine, M., Kosugi, H., Matsuo, Y., Fujisawa, T., Omata, S., Kishi, E., Shimizu, A., Tsukatani, N., Tanikawa, S., Fujita, N., & Harayama, S. (2008). Complete genome sequence of the soil actinomycete *Kocuria rhizophila*. Journal of Bacteriology, 190(12), 4139–4146. 10.1128/JB.01853-07

Vassallo, L., Hansmann, D., & Braunstein, L. A. (2019). On the growth of non-motile bacteria colonies: an agent-based model for pattern formation. European Physical Journal B, 92(9). 10.1140/epjb/e2019-100265-0

Wickham, H., François, R., Henry, L., & Müller, K. (2023). dplyr: A Grammar of Data Manipulation. R package. https://github.com/tidyverse/dplyr, https://dplyr.tidyverse.org.

Wickham H. (2016). ggplot2: Elegant Graphics for Data Analysis. Springer-Verlag New York, ISBN 978-3-319-24277-4. https://ggplot2.tidyverse.org.

Xue, H., Kurokawa, M., & Ying, B. W. (2021). Correlation between the spatial distribution and colony size was common for monogenetic bacteria in laboratory conditions. BMC Microbiology, 21(1), 1–9. 10.1186/s12866-021-02180-8

Ye, Y., Ghrayeb, M., Miercke, S., Arif, S., Müller, S., Mascher, T., Chai, L., & Zaburdaev, V. (2024) Residual cells and nutrient availability guide wound healing in bacterial biofilms. Soft Matter, 20, 1047–1060. doi: 10.1039/d3sm01032e. PMID: 38205608.

Yordanov, S., Neuhaus, K., Hartmann, R., Díaz-Pascual, F., Vidakovic, L., Singh, P. K., & Drescher, K. (2021). Single-objective high-resolution confocal light sheet fluorescence microscopy for standard biological sample geometries. Biomedical Optics Express, 12(6), 3372. 10.1364/boe.420788

He, Z., Kisla, D., Zhang, L., Yuan, C., Green-Church, K.B., Yousef, A.E. 2007. Isolation and identification of a *Paenibacillus polymyxa* strain that coproduces a novel lantibiotic and polymyxin. Appl Environ Microbiol 73: 10.1128/AEM.02023-06

Zhang, M., Zhang, J., Wang, Y., Wang, J., Achimovich, A. M., Acton, S. T., & Gahlmann, A. (2020). Non-invasive single-cell morphometry in living bacterial biofilms. Nature Communications, 11(1), 1–13. 10.1038/s41467-020-19866-8

Zheng, S., Bawazir, M., Dhall, A., Kim, H. E., He, L., Heo, J., & Hwang, G. (2021). Implication of surface properties, bacterial motility, and hydrodynamic conditions on bacterial surface sensing and their initial adhesion. Frontiers in Bioengineering and Biotechnology, 9(February), 1–22. 10.3389/fbioe.2021.643722

