## Supplementary file for "Bacterial colony biopsies: spatial discrimination of heterogeneous cell types by cytometric fingerprinting"

#### Supplementary Information

### Cytometric biopsies: high-resolution discrimination of heterogeneous cell types spatially distributed in clonal microbial colonies

Gorkhmaz Abbaszade, Kathrin Stückrath and Susann Müller

Department of Applied Microbial Ecology, Helmholtz-Centre for Environmental Research - UFZ, Permoserstr. 15, 04318 Leipzig, Germany

Corresponding author
Susann Müller, Department of Applied Microbial Ecology, Helmholtz-Centre for Environmental Research - UFZ, Permoserstr. 15, 04318 Leipzig, Germany,

##### Content

SI Figure 1: Images of colonies of the strains *Bacillus subtilis*, *Paenibacillus polymyxa*, *Kocuria rhizophila*, *Stenotrophomonas rhizophila*, and *Pseudomonas citronellolis*.

SI Figure 2: Flow cytometric analysis of biopsy samples from colonies of *Bacillus subtilis* (FSC vs. DAPI).
SI Figure 3: Flow cytometric analysis of biopsy samples from colonies of *Paenibacillus polymyxa* (FSC vs. DAPI).
SI Figure 4: Flow cytometric analysis of biopsy samples from colonies of *Kocuria rhizophila* (FSC vs. DAPI).
SI Figure 5: Flow cytometric analysis of biopsy samples from colonies of *Stenotrophomonas rhizophila* (FSC vs. DAPI).
SI Figure 6: Flow cytometric analysis of biopsy samples from colonies of *Pseudomonas citronellolis* (FSC vs. DAPI)

SI Figure 7: Flow cytometric analysis of biopsy samples from colonies of *Bacillus subtilis* (SYTO9 vs. PI). SI Figure 8: Flow cytometric analysis of biopsy samples from colonies of *Paenibacillus polymyxa* (SYTO9 vs. PI).
SI Figure 9: Flow cytometric analysis of biopsy samples from colonies of *Kocuria rhizophila* (SYTO9 vs. PI).
SI Figure 10: Flow cytometric analysis of biopsy samples from colonies of *Stenotrophomonas rhizophila* (SYTO9 vs. PI).
SI Figure 11: Flow cytometric analysis of biopsy samples from colonies of *Pseudomoas citronellolis* (SYTO9 vs. PI).

SI Figure 12: Comparison of different fixation procedures for the strains *Bacillus subtilis*, *Paenibacillus polymyxa*, *Kocuria rhizophila*, *Stenotrophomonas rhizophila* and *Pseudomonas citronellolis*.

SI Figure 13: Similarity comparison of biopsy sites sampled from a 3-day-old *Paenibacillus polymyxa* colony using different needle or tip sizes.
SI Figure 14: Similarity comparison of biopsy sites sampled from a 3-day-old *Kocuria rhizophila* colony using different needle or tip sizes.
SI Figure 15: Similarity comparison of biopsy sites sampled from a 3-day-old *Stenotrophomonas rhizophila* colony using different needle or tip sizes.
SI Figure 16: Similarity comparison of biopsy sites sampled from a 3-day-old *Pseudomonas citronellolis* colony using different needle or tip sizes.

SI Figure 17: Proportions of cell types in *Bacillus subtilis* colonies.
SI Figure 18: Proportions of cell types in *Paenibacillus polymyxa* colonies.
SI Figure 19: Proportions of cell types in *Kocuria rhizophila* colonies.
SI Figure 20: Proportions of cell types in *Stenotrophomonas rhizophila* colonies.
SI Figure 21: Proportions of cell types in *Pseudomonas citronellolis* colonies.

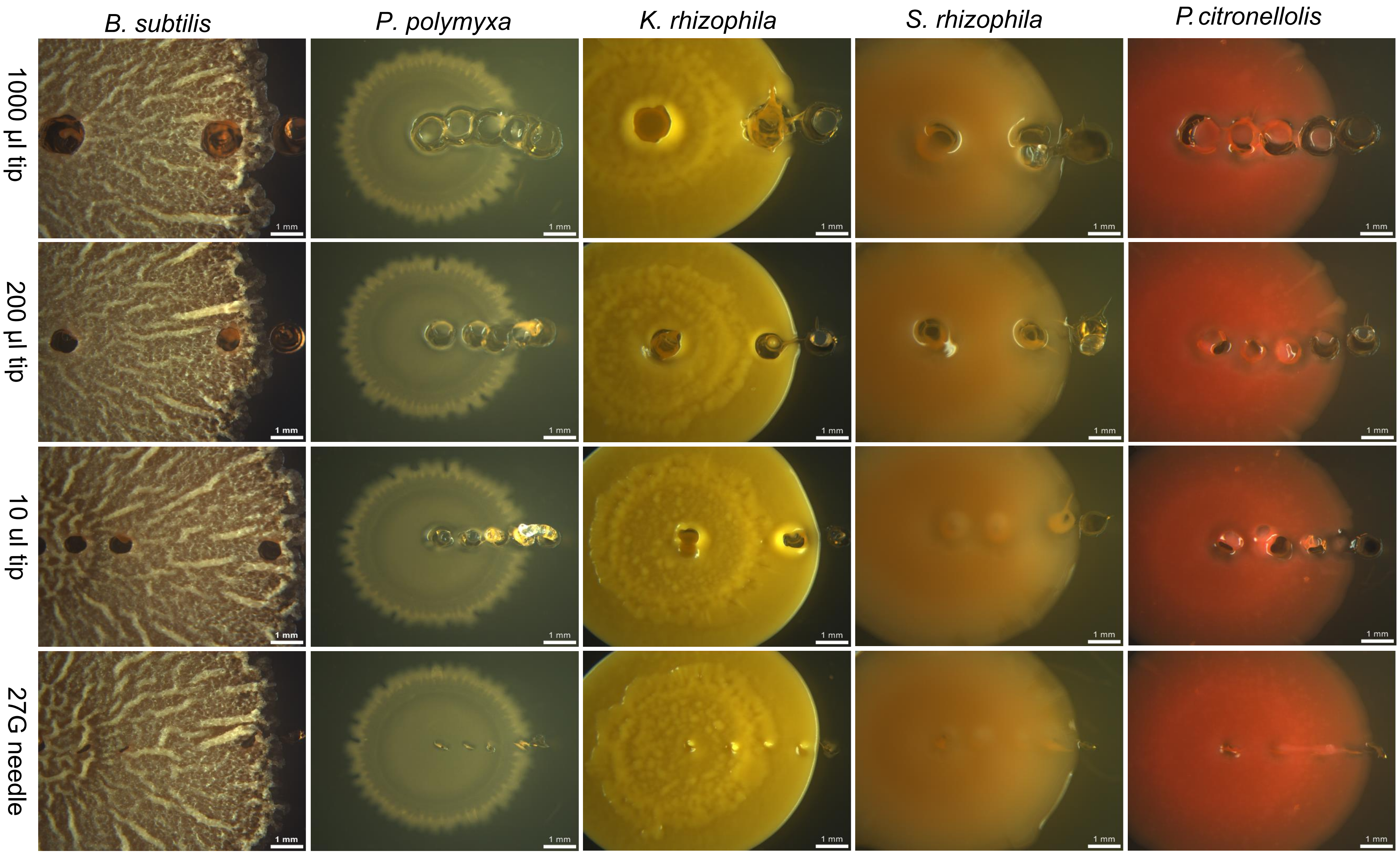


**SI Figure 1**: Images of colonies of the strains *Bacillus subtilis*, *Paenibacillus polymyxa*, *Kocuria rhizophila*, *Stenotrophomonas rhizophila*, and *Pseudomonas citronellolis*. The locations of the biopsies are shown and also the differences between 27G needling and using tips of sizes 10 µL – 1000 µL. The colonies were all grown for 3 days. The locations of the biopsies are summarized in Figure 1.

#### *Bacillus subtilis* (NCIB 3610)

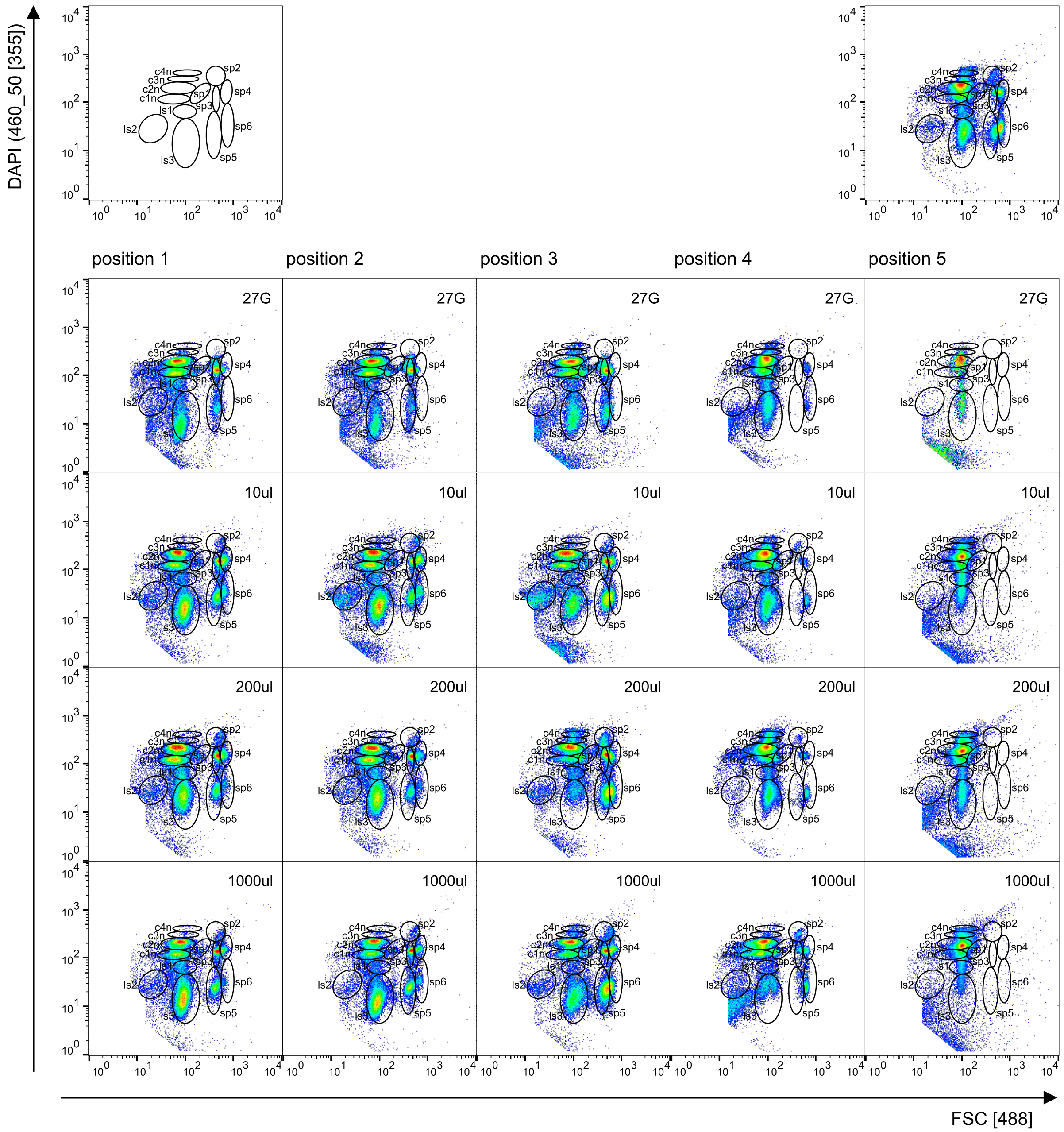


**SI Figure 2**: Flow cytometric analysis of biopsy samples from 3-day-old colonies of *Bacillus subtilis*. Colonies were biopsied using a 27G needle and 10 µL to 1000 µL tips. The cells were fixated and stained as described in the methods. A cell gate separated 50.000 cells from debris and instrumental noise (not shown). The cells were measured with regard to FSC vs. DAPI fluorescence. Left above: gate template highlighting 4 proliferating cell subgroups (c1n-c4n), 6 spore types (sp1-sp6) and 3 less stained cell types (ls1-ls3). The proliferating cell subgroups were determined according to the DAPI fluorescence intensity values on the y-axis with c1n-c4n marking doublings in fluorescence intensities. Right above: analysis of the whole colony. Middle: Position of the biopsies in the colonies and type of needle and tips used. The numbers of the cells in the gates were the basis for creating the Figures 3 and 5 - 7 in the main manuscript and the SI Figures 13 - 21.

#### *Paenibacillus polymyxa* (DSM 36)

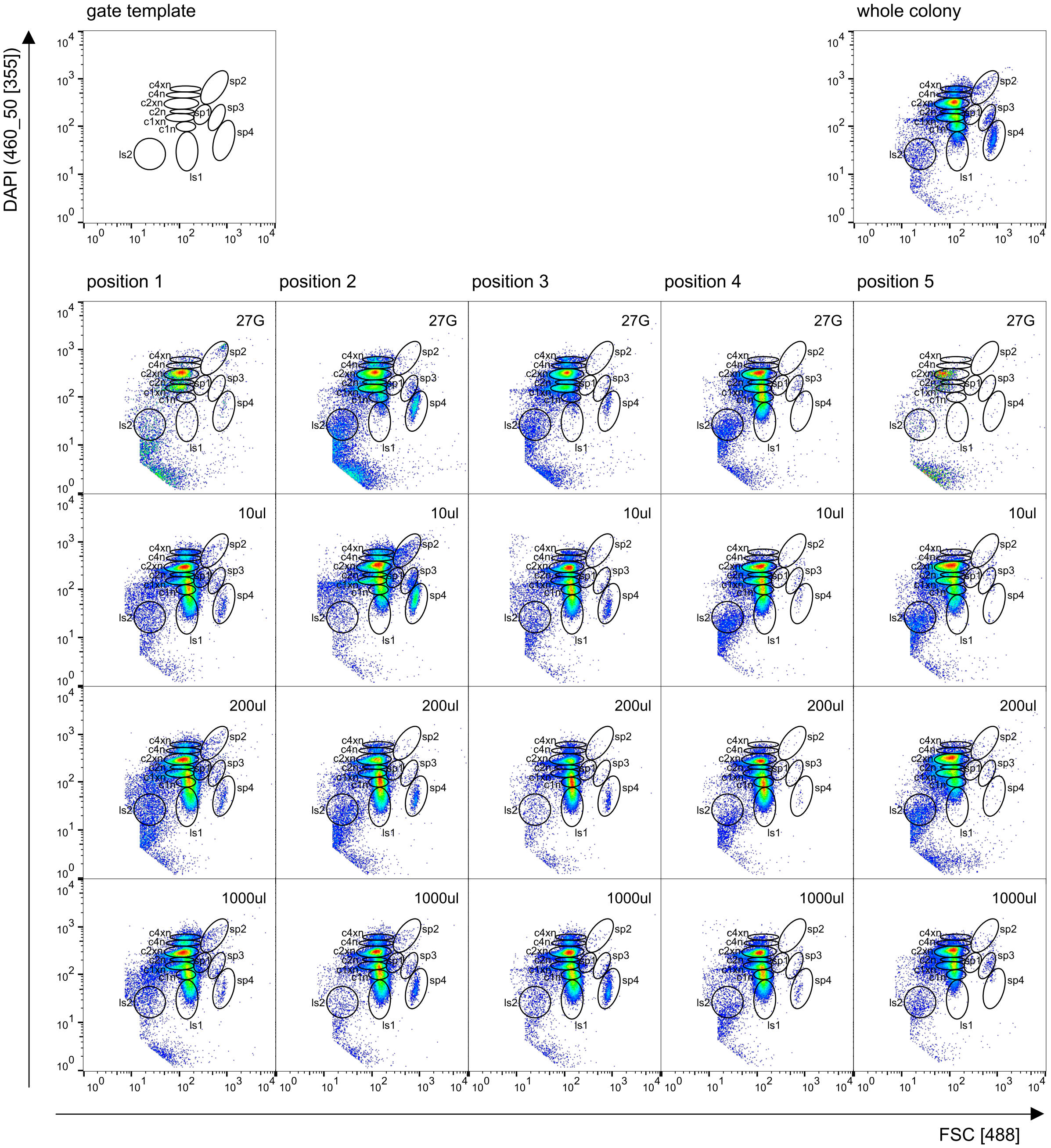


**SI Figure 3**: Flow cytometric analysis of biopsy samples from 3-day-old colonies of *Paenibacillus polymyxa*. Colonies were biopsied using a 27G needle and 10 µL to 1000 µL tips. The cells were fixated and stained as described in the methods. A cell gate separated 50.000 cells from debris and instrumental noise (not shown). The cells were measured with regard to FSC vs. DAPI fluorescence. Left above: gate template highlighting 6 proliferating cell subgroups (c1n-c4xn), 4 spore types (sp1-sp4) and 2 less stained cell types (ls1, ls2). The proliferating cell subgroups were determined according to the DAPI fluorescence intensity values on the y-axis with c1n-c4n marking doublings in fluorescence intensities and c1xn-c4xn DAPI intensities in-between, highlighting transition states in proliferation. Right above: analysis of the whole colony. Middle: Position of the biopsies in the colonies and type of needle and tips used. The numbers of the cells in the gates were the basis for creating the Figures 3 and 5 - 7 in the main manuscript and the SI Figures 13 - 21.

#### *Kocuria rhizophila* (DSM 348)

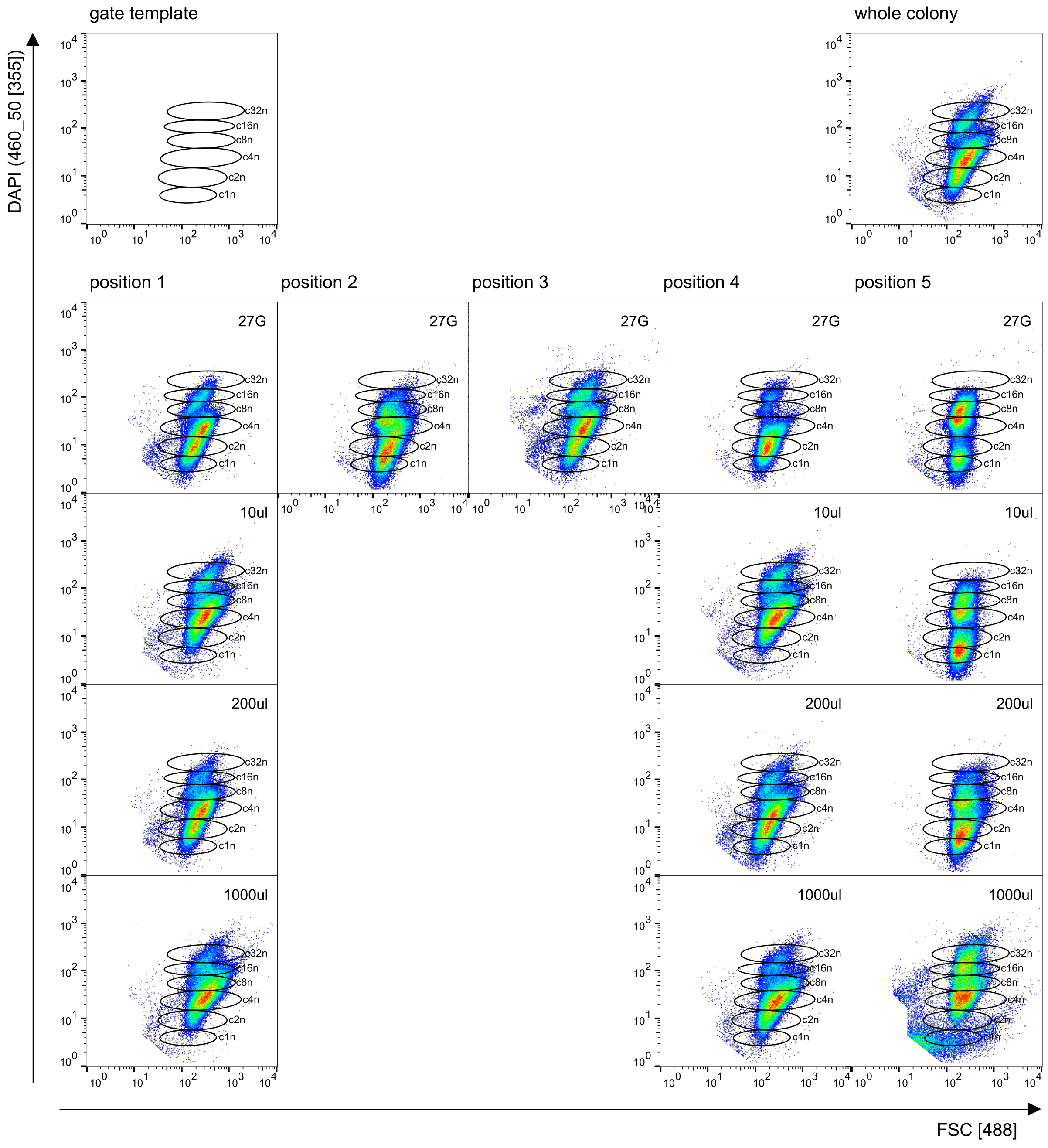


**SI Figure 4**: Flow cytometric analysis of biopsy samples from 3-day-old colonies of *Kocuria rhizophila*. Colonies were biopsied using a 27G needle and 10 µL to 1000 µL tips. The cells were fixated and stained as described in the methods. A cell gate separated 50.000 cells from debris and instrumental noise (not shown). The cells were measured with regard to FSC vs. DAPI fluorescence. Left above: gate template highlighting 6 proliferating cell subgroups (c1n-c32n). The proliferating cell subgroups were determined according to the DAPI fluorescence intensity values on the y-axis with c1n-c32n marking doublings in fluorescence intensities. Right above: analysis of the whole colony. Middle: Position of the biopsies in the colonies and type of needle and tips used. The numbers of the cells in the gates were the basis for creating the Figures 3 and 5 - 7 in the main manuscript and the SI Figures 13 - 21.

#### *Stenotrophomonas rhizophila* (DSM 14405)

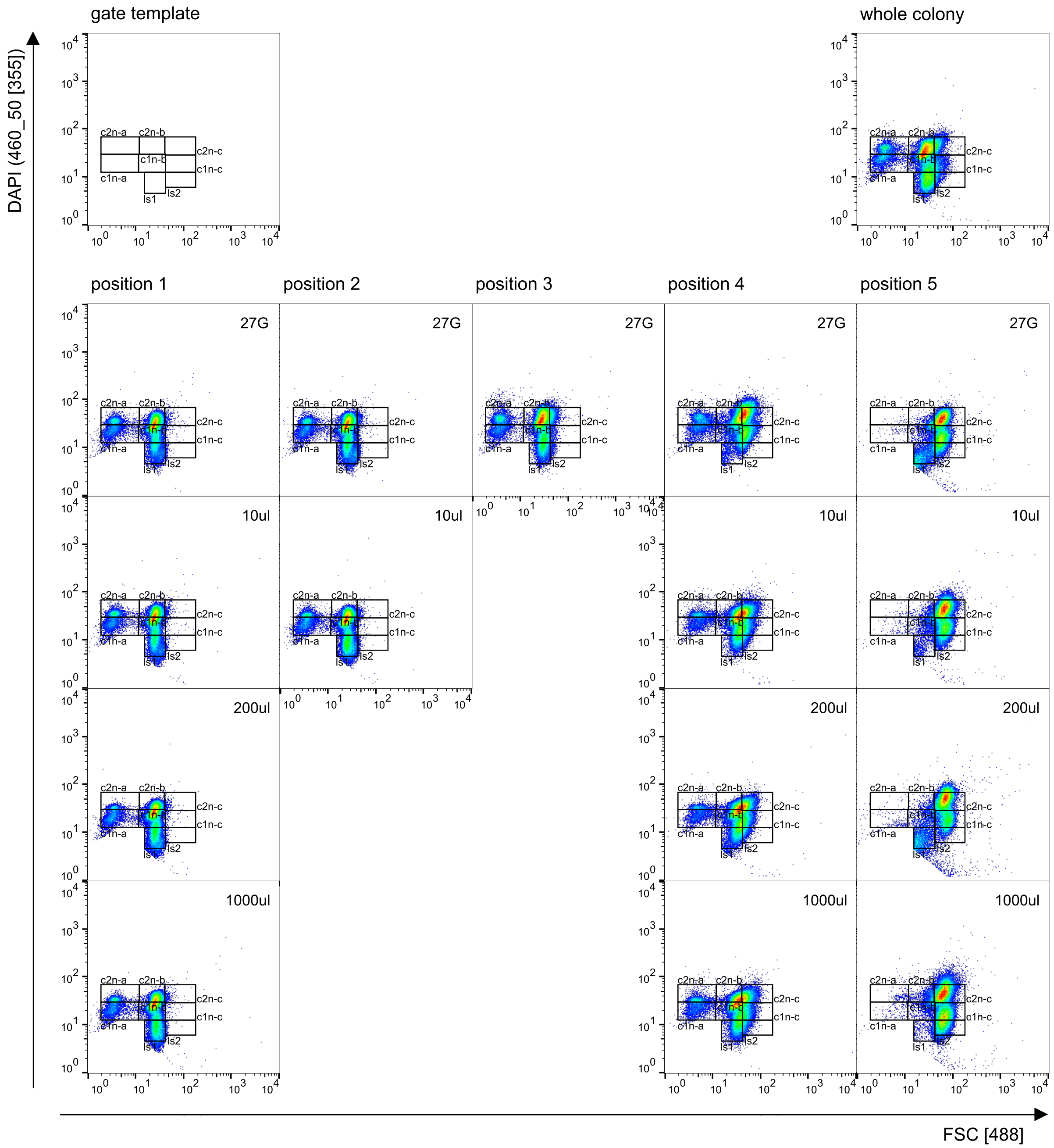


**SI Figure 5:** Flow cytometric analysis of biopsy samples from 3-day-old colonies of *Stenotrophomonas rhizophila*. Colonies were biopsied using a 27G needle and 10 µL to 1000 µL tips. The cells were fixated and stained as described in the methods. A cell gate separated 50.000 cells from debris and instrumental noise (not shown). The cells were measured with regard to FSC vs. DAPI fluorescence. Left above: gate template highlighting 6 proliferating cell subgroups (c1n:a-c, c2n:a-c) and 2 less stained cell types (ls1, ls2). The proliferating cell subgroups were determined according to the DAPI fluorescence intensity values on the y-axis with c1n-c2n marking doublings in fluorescence intensities. The strain shows 2 subgroups of cells with regard to cell size (a-c). Right above: analysis of the whole colony. Middle: Position of the biopsies in the colonies and type of needle and tips used. The numbers of the cells in the gates were the basis for creating the Figures 3 and 5 - 7 in the main manuscript and the SI Figures 13 - 21.

#### *Pseudomonas citronellolis* (P3B5)

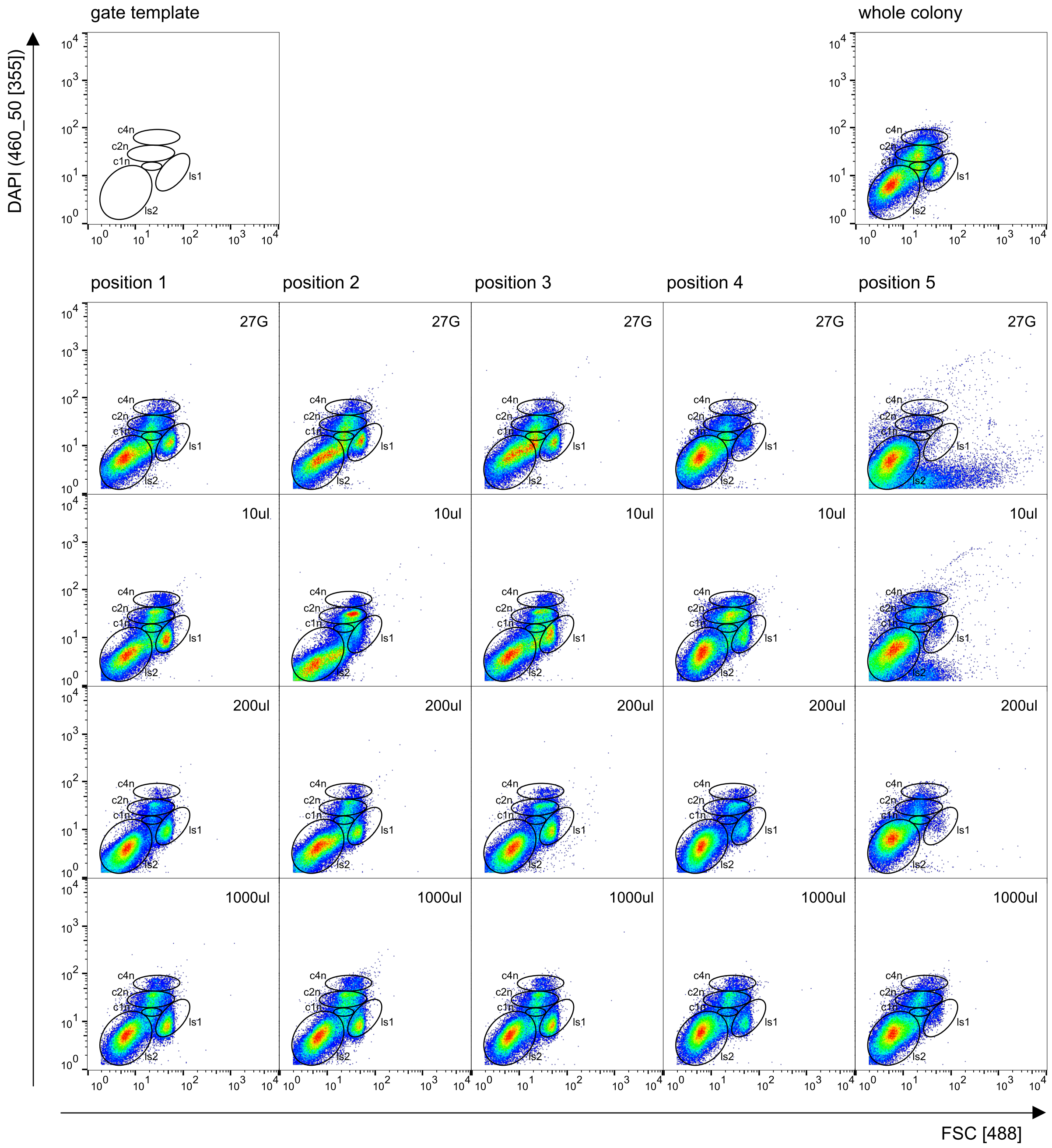


**SI Figure 6**: Flow cytometric analysis of biopsy samples from 3-day-old colonies of *Pseudomonas citronellolis*. Colonies were biopsied using a 27G needle and 10 µL to 1000 µL tips. The cells were fixated and stained as described in the methods. A cell gate separated 50.000 cells from debris and instrumental noise (not shown).The cells were measured with regard to FSC vs. DAPI fluorescence. Left above: gate template highlighting 3 proliferating cell subgroups (c1n-c4n) and 2 less stained cell types (ls1, ls2). The proliferating cell subgroups were determined according to the DAPI fluorescence intensity values on the y-axis with c1n-c2n marking doublings in fluorescence intensities. Right above: analysis of the whole colony. Middle: Position of the biopsies in the colonies and type of needle and tips used. The numbers of the cells in the gates were the basis for creating the Figures 3 and 5 - 7 in the main manuscript and the SI Figures 13 - 21.

#### ***Bacillus subtilis* (NCIB 3610)**

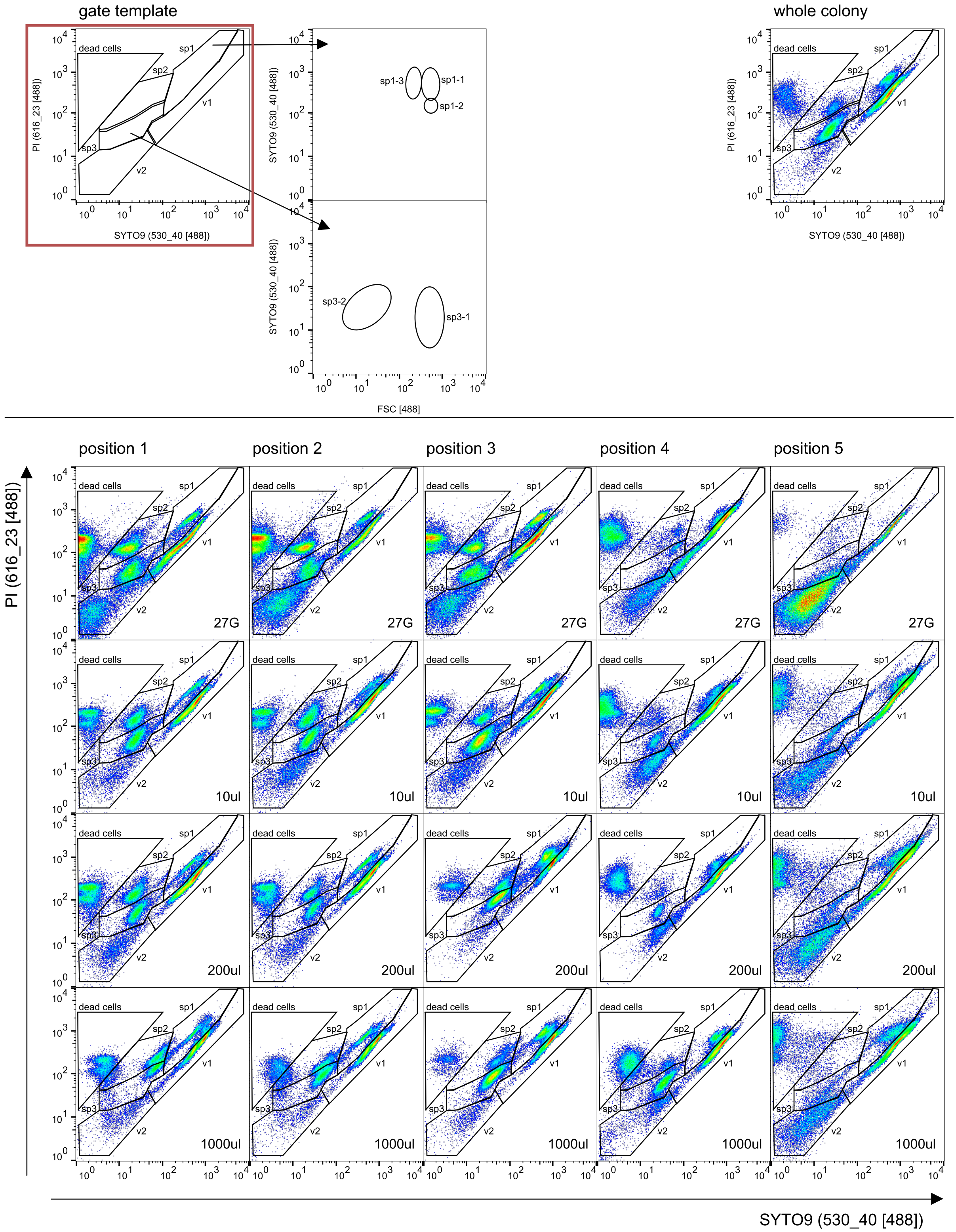


**SI Figure 7**: Flow cytometric analysis of biopsy samples from 3-day-old colonies of *Bacillus subtilis*. Colonies were biopsied using a 27G needle and 10 µL to 1000 µL tips. The cells were live stained as described in the methods. A cell gate separated 50.000 cells from debris and instrumental noise (not shown). The cells were measured with regard to SYTO9 vs. PI fluorescence. Left above: gate template highlighting 2 vegetative cell subgroups (v1-v2), 1 dead cell group and 3 spore types (sp1-sp3). The spore types sp1 and sp3 further differentiate into 3 and 2 subgroups in the FSC, respectively (not shown), therefore representing 6 different spore types. The vegetative cells and spores were discriminated by SYTO9 fluorescence while PI marks membrane and cell wall compromised cells. Right above: analysis of the whole colony. Middle: Position of the biopsies in the colonies and type of needle and tips used. The numbers of the cells in the gates were the basis for creating the Figures 3 and 5 - 7 in the main manuscript and the SI Figures 13 - 21.

#### *Paenibacillus polymyxa* (DSM 36)

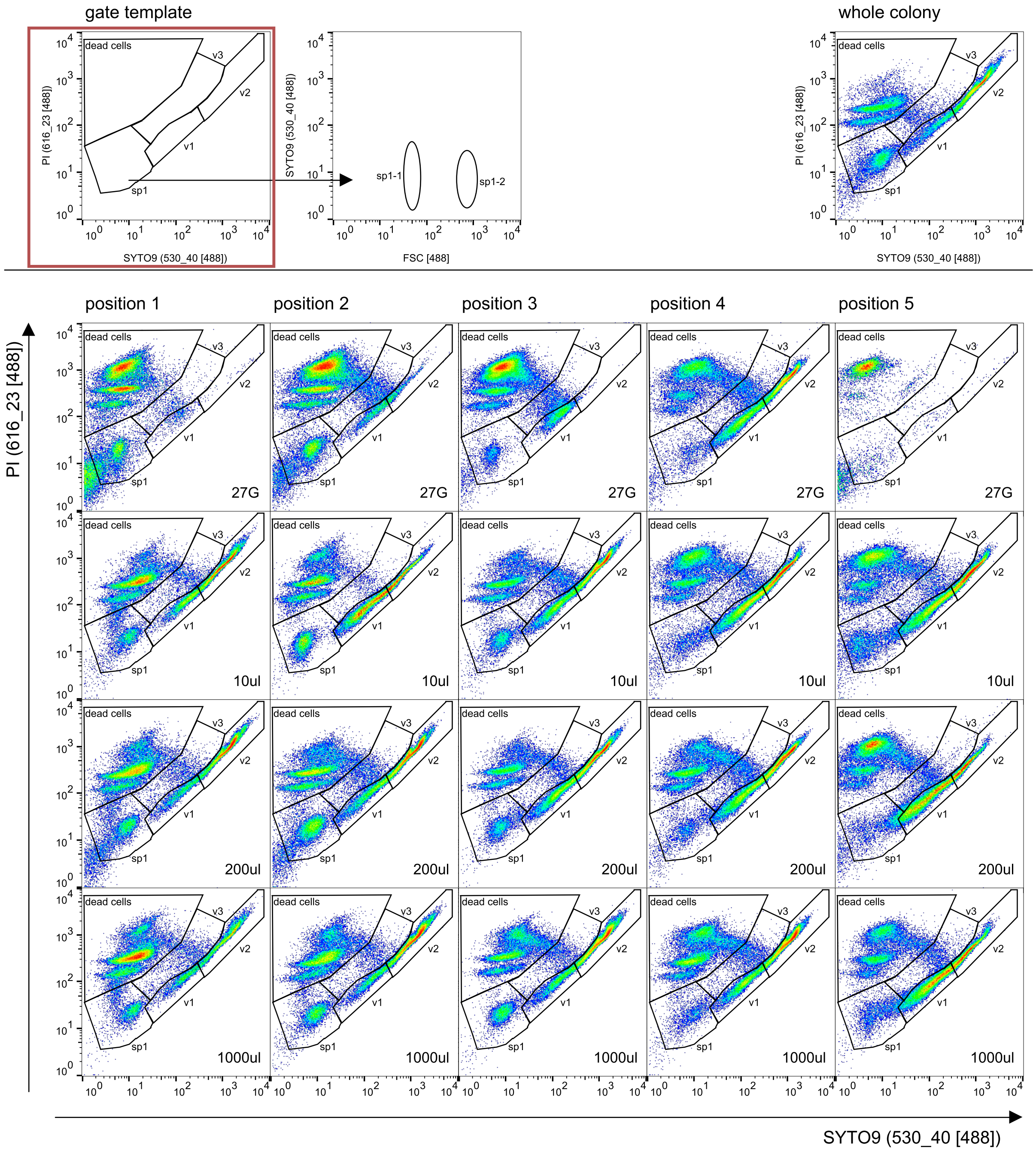


**SI Figure 8:** Flow cytometric analysis of biopsy samples from 3-day-old colonies of *Paenibacillus polymyxa*. Colonies were biopsied using a 27G needle and 10 µL to 1000 µL tips. The cells were live stained as described in the methods. A cell gate separated 50.000 cells from debris and instrumental noise (not shown). The cells were measured with regard to SYTO9 vs. PI fluorescence. Left above: gate template highlighting 3 vegetative cell subgroups (v1-v3), 1 dead cell group and 1 spore type (sp1). The spore type sp1 further differentiate into 2 subgroups in the FSC (not shown), therefore representing 2 different spore types. The vegetative cells and spores were discriminated by SYTO9 fluorescence while PI marks membrane and cell wall compromised cells. Right above: analysis of the whole colony. Middle: Position of the biopsies in the colonies and type of needle and tips used. The numbers of the cells in the gates were the basis for creating the Figures 3 and 5 - 7 in the main manuscript and the SI Figures 13 - 21.

***Kocuria rhizophila* (DSM 348)**

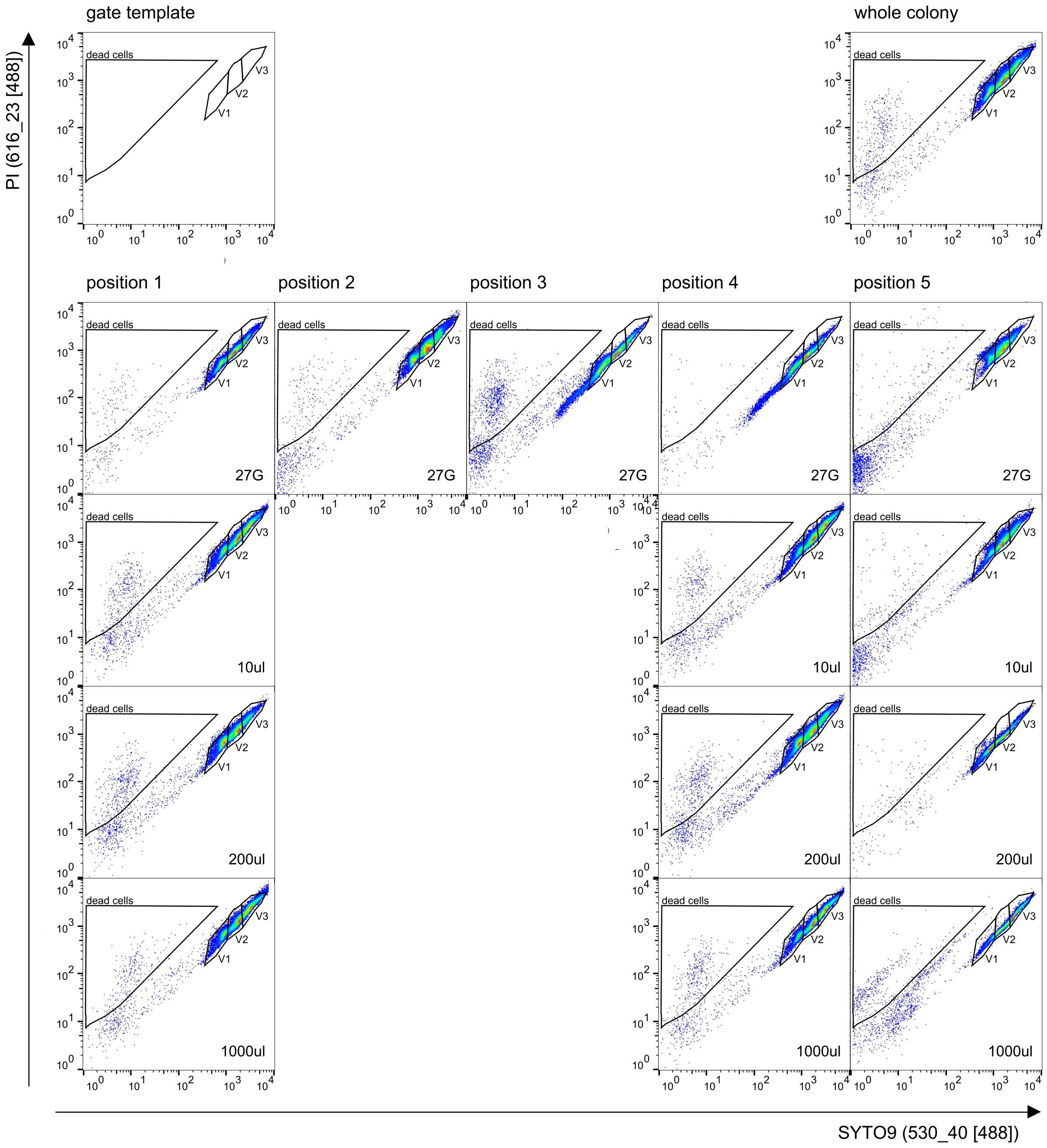


**SI Figure 9**: Flow cytometric analysis of biopsy samples from 3-day-old colonies of *Kocuria rhizophila*. Colonies were biopsied using a 27G needle and 10 µL to 1000 µL tips. The cells were live stained as described in the methods. A cell gate separated 50.000 cells from debris and instrumental noise (not shown). The cells were measured with regard to SYTO9 vs. PI fluorescence. Left above: gate template highlighting 3 vegetative cell subgroups (v1-v3) and 1 dead cell group. The vegetative cells were discriminated by SYTO9 fluorescence while PI marks membrane and cell wall compromised cells. Right above: analysis of the whole colony. Middle: Position of the biopsies in the colonies and type of needle and tips used. The numbers of the cells in the gates were the basis for creating the Figures 3 and 5 - 7 in the main manuscript and the SI Figures 13 - 21.

#### *Stenotrophomonas rhizophila* (DSM 14405)

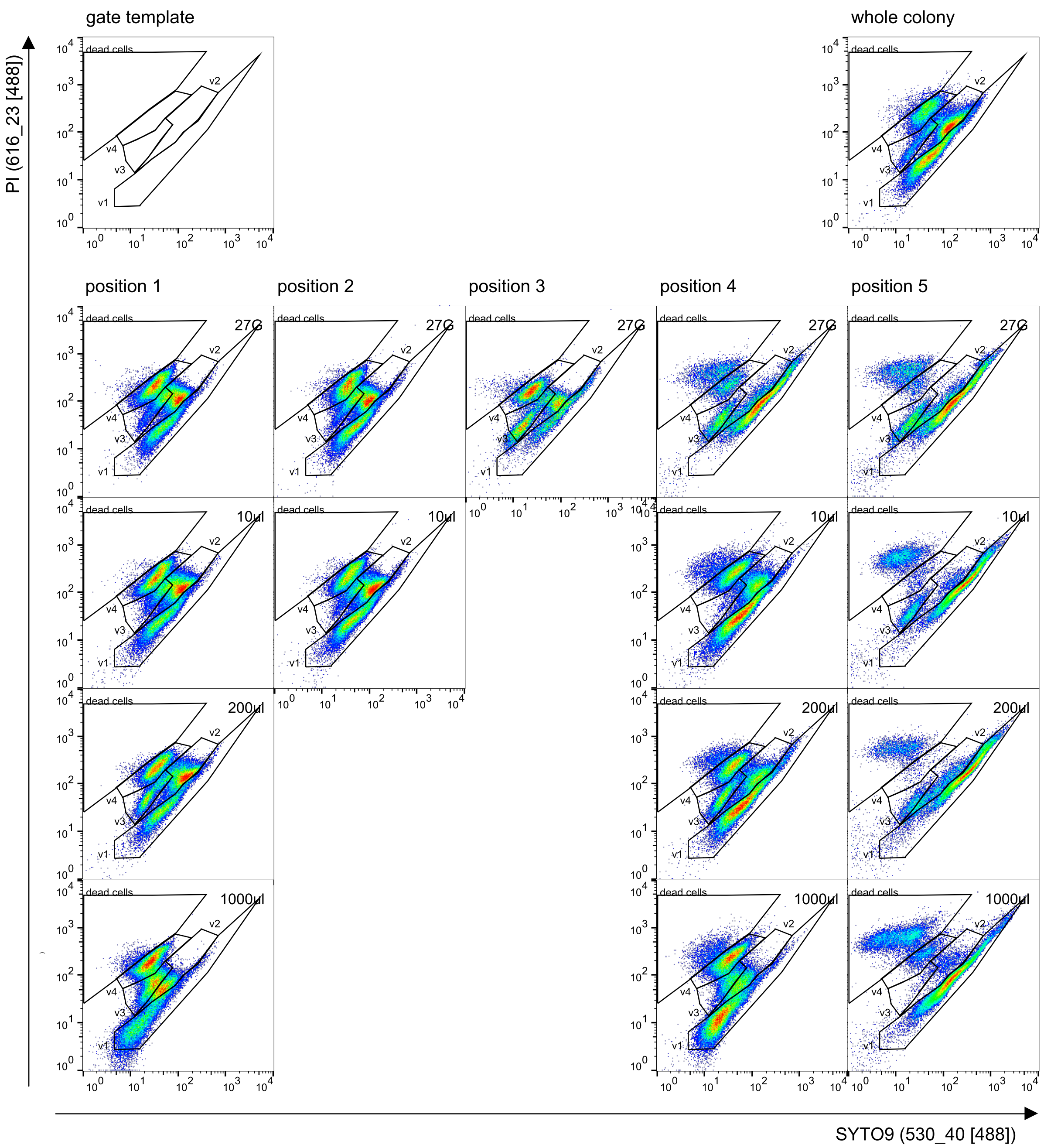


**SI Figure 10**: Flow cytometric analysis of biopsy samples from 3-day-old colonies of *Stenotrophomonas rhizophila*. Colonies were biopsied using a 27G needle and 10 µL to 1000 µL tips. The cells were live stained as described in the methods. A cell gate separated 50.000 cells from debris and instrumental noise (not shown). The cells were measured with regard to SYTO9 vs. PI fluorescence. Left above: gate template highlighting 4 vegetative cell subgroups (v1-v4), and 1 dead cell group. The vegetative cells and less stained cell types were discriminated by SYTO9 fluorescence while PI marks membrane and cell wall compromised cells. Right above: analysis of the whole colony. Middle: Position of the biopsies in the colonies and type of needle and tips used. The numbers of the cells in the gates were the basis for creating the Figures 3 and 5 - 7 in the main manuscript and the SI Figures 13 - 21.

#### *Pseudomonas citronellolis* (P3B5)

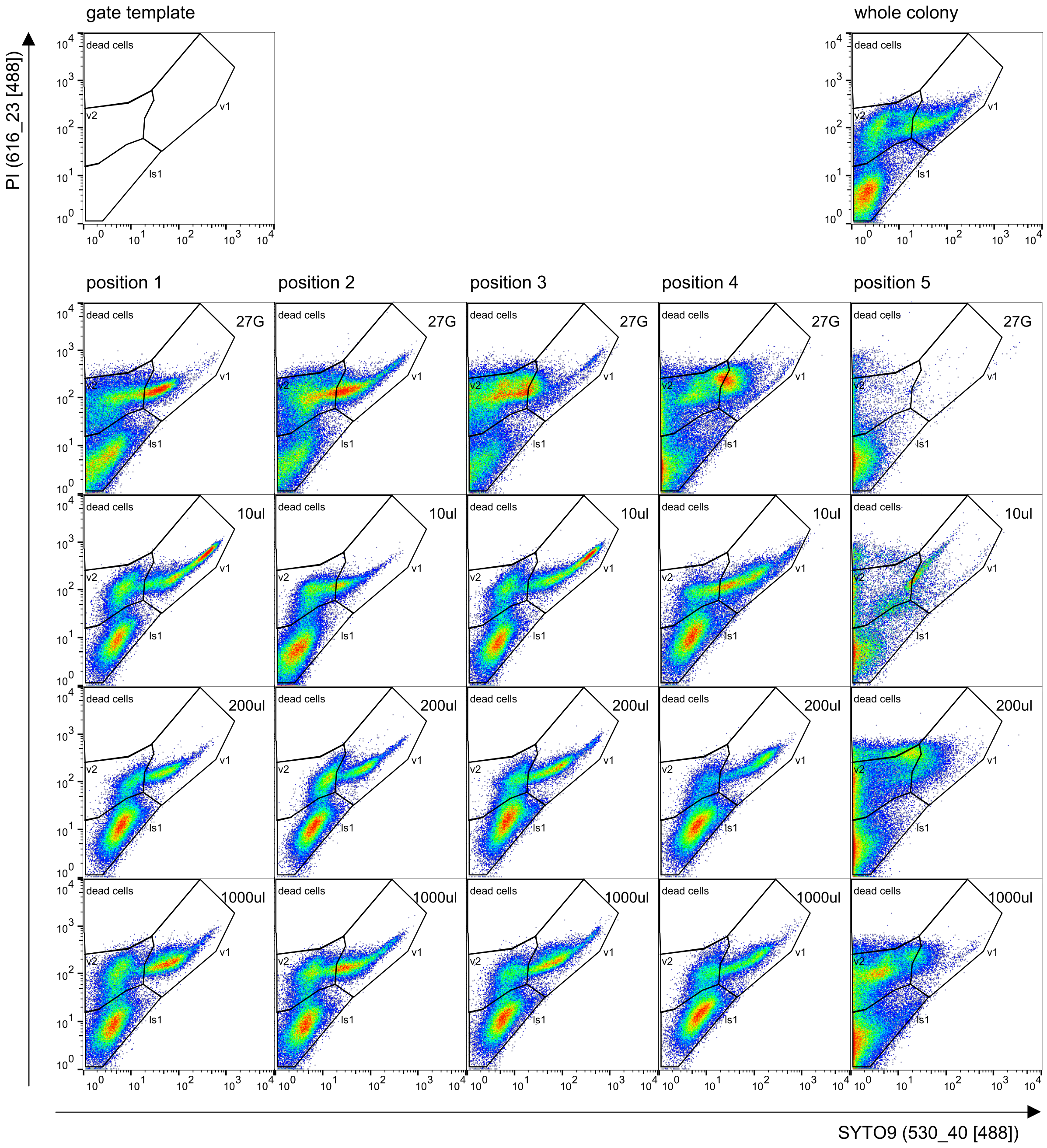


**SI Figure 11**: Flow cytometric analysis of biopsy samples from 3-day-old colonies of *Pseudomoas citronellolis*. Colonies were biopsied using a 27G needle and 10 µL to 1000 µL tips. The cells were live stained as described in the methods. A cell gate separated 50.000 cells from debris and instrumental noise (not shown). The cells were measured with regard to SYTO9 vs. PI fluorescence. Left above: gate template highlighting 2 vegetative cell subgroups (v1-v2), 1 dead cell group and 1 less stained events (ls1). The vegetative cells and less stained events were discriminated by SYTO9 fluorescence while PI marks membrane and cell wall compromised cells. Please be aware that this strain is heavily pumping SYTO9 and therefore the numbers of cells in gates are constantly changing directly during measurement. Right above: analysis of the whole colony. Middle: Position of the biopsies in the colonies and type of needle and tips used. The numbers of the cells in the gates were the basis for creating the Figures 3 and 5 - 7 in the main manuscript and the SI Figures 13 - 21.

#### *Bacillus subtilis* (NCIB 3610)

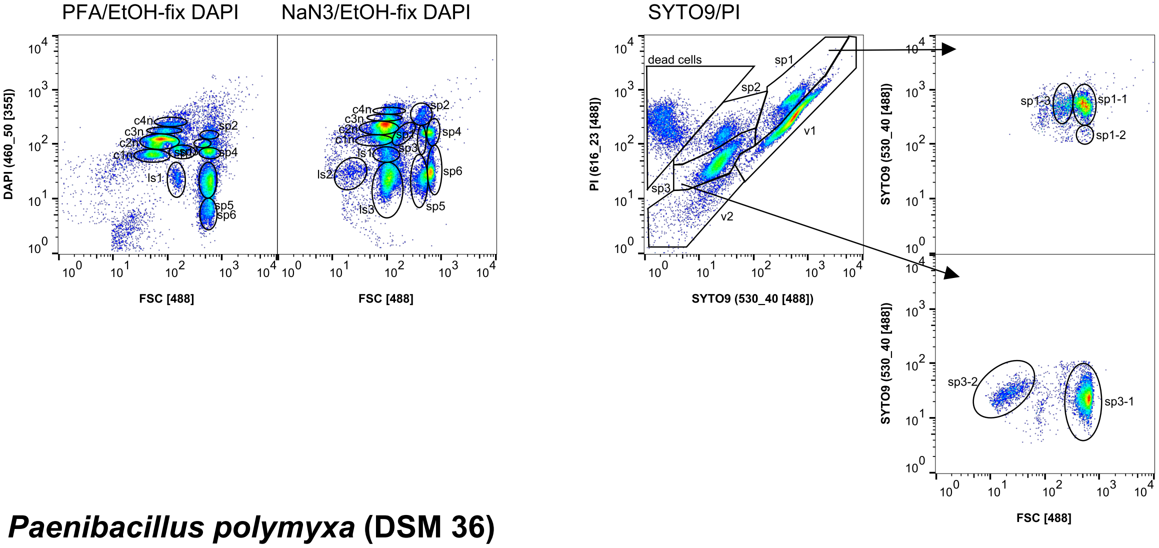


#### *Paenibacillus polymyxa* (DSM 36)

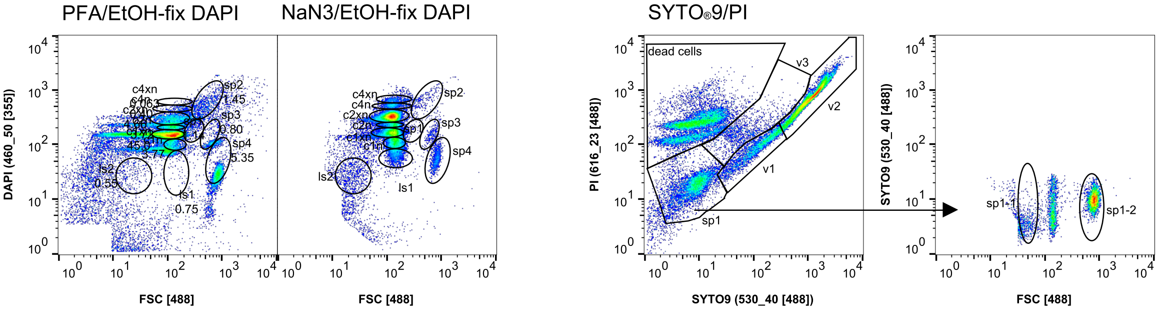


#### *Kocuria rhizophila* (DSM 348)

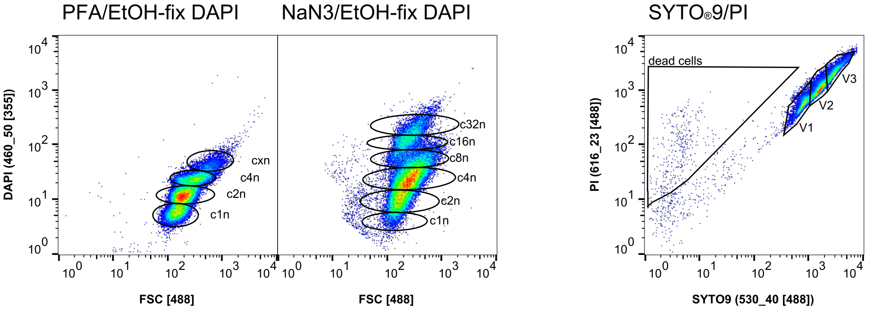


#### *Stenotrophomonas rhizophila* (DSM 14405)

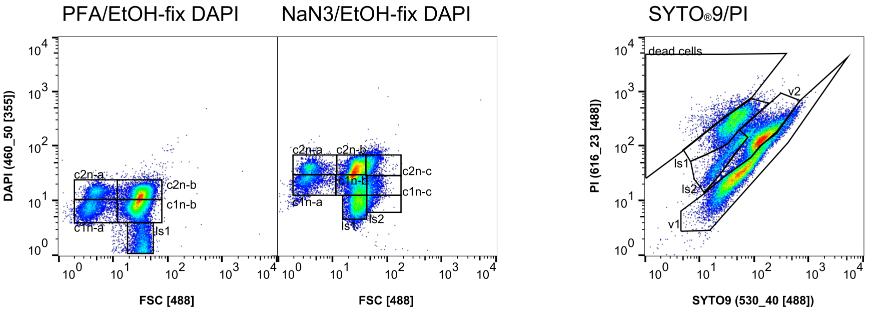

#### ***Pseudomonas citronellolis* (P3B5)**

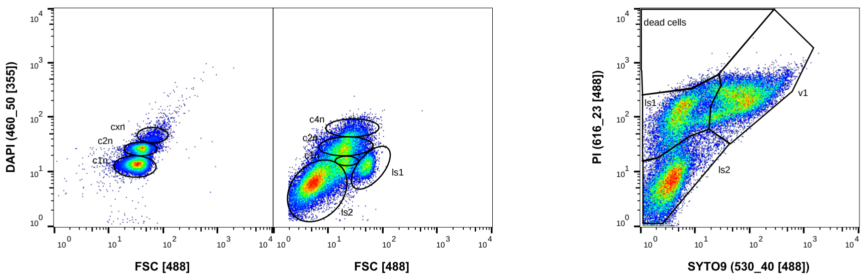


**SI Figure 12**: Comparison of different fixation procedures for the strains *Bacillus subtilis*, *Paenibacillus polymyxa*, *Kocuria rhizophila*, *Stenotrophomonas rhizophila*, and *Pseudomonas citronellolis*. Left: standard PFA/EtOH fixation with following DAPI staining, Middle: $NaN_3$/EtOH fixation with following DAPI staining. Right: live cell measurement with SYTO9 and PI staining plus back gating for FSC vs. SYTO9 in case of *Bacillus subtilis* and *Paenibacillus polymyxa*.

**_Paenibacillus polymyxa_ (DSM 36)**

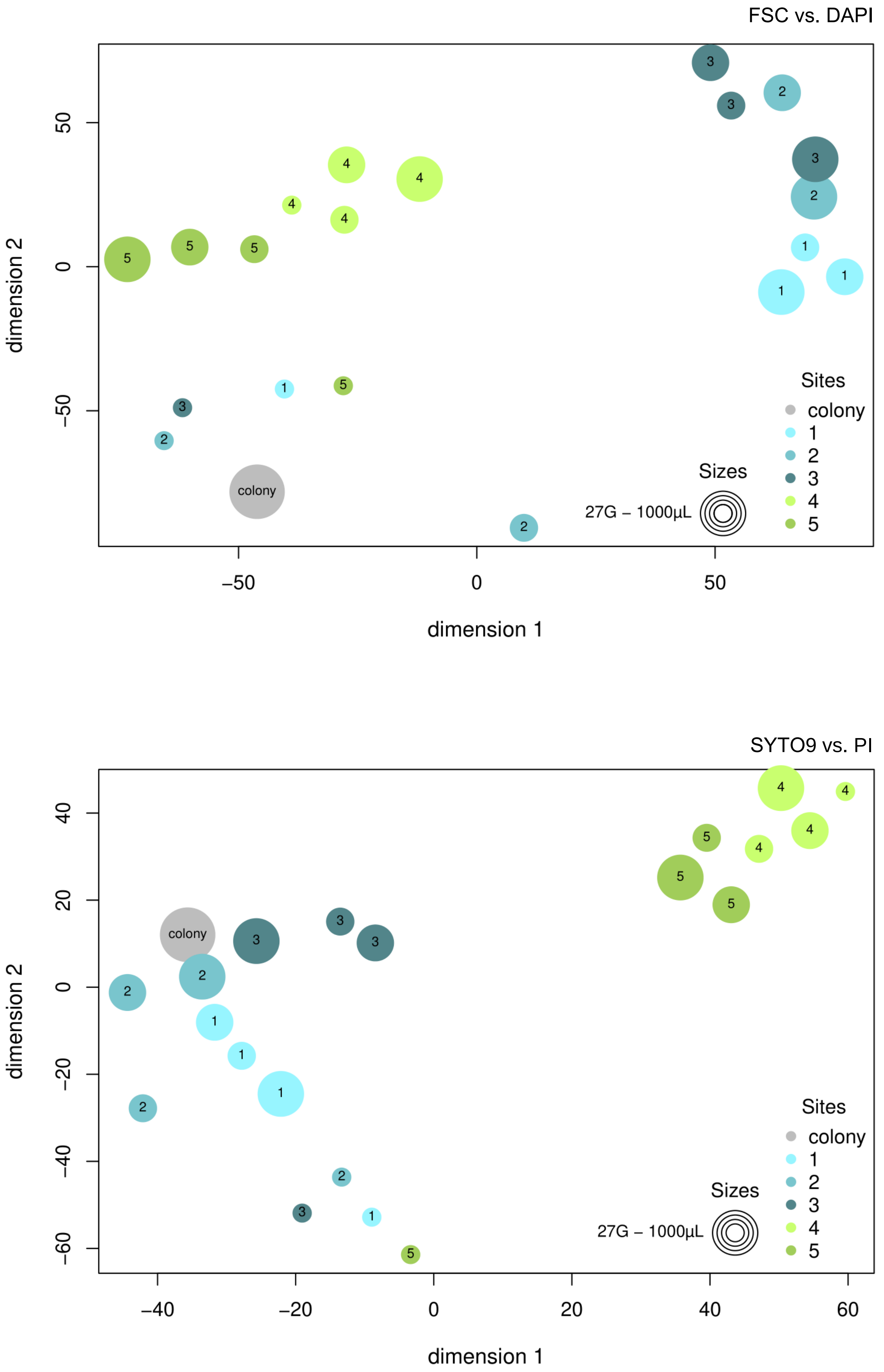


**SI Figure 13**: Similarity comparison of biopsy sites sampled from a 3-day-old *Paenibacillus polymyxa* colony using different needle or tip sizes. Samples were collected from five distinct positions within the colony (labeled 1 to 5 in respective colors, as detailed in Table 2), and measured by flow cytometry to create cytometric fingerprints (2D plots). The size of the points reflects the dimensions of the sampling tips: 27G needle, as well as 10 µL, 200 µL and 1000 µL pipette tips. Grey point: whole colony. Above: Samples were fixated and stained by the DAPI staining method. Below: Live samples were stained with the SYTO9 and PI staining method. The cell numbers in gates of the 2D plots were transformed by the Box-Cox transformation method and visualized by the R package tSNE.

***Kocuria rhizophila* (DSM 348)**

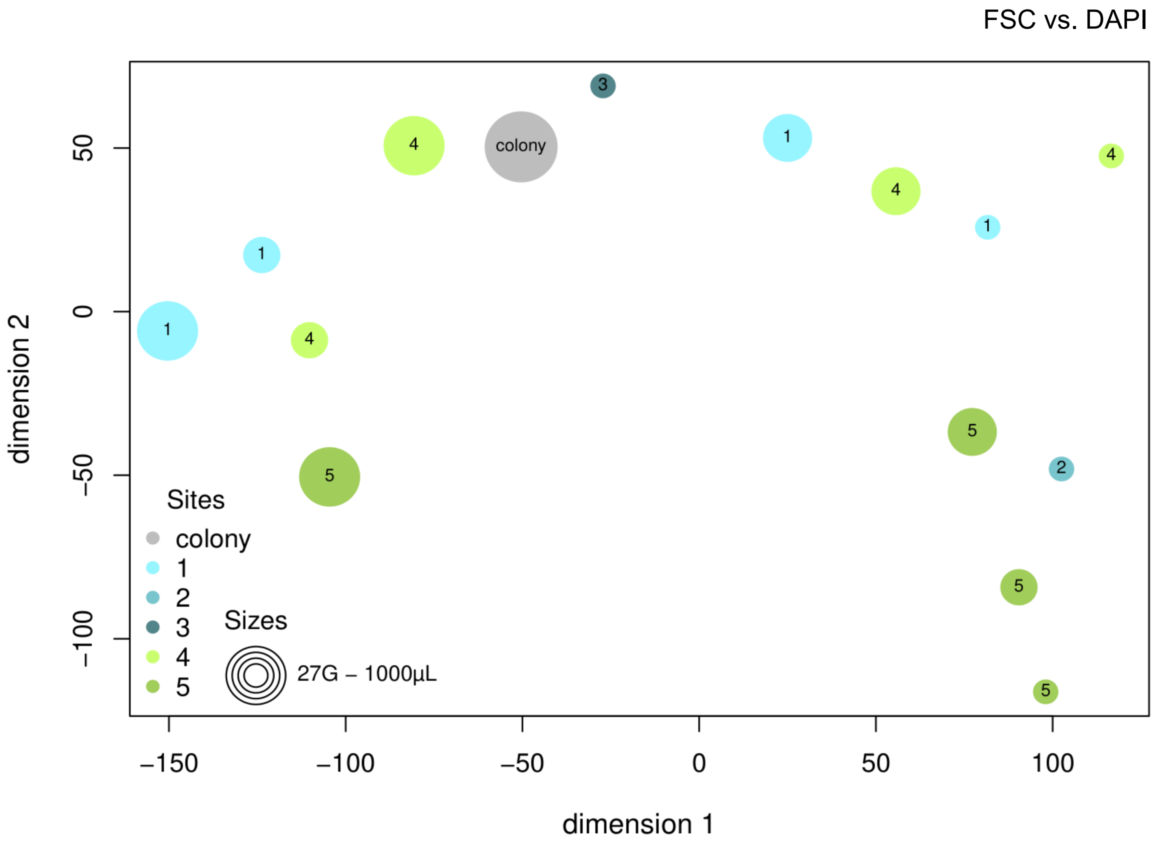


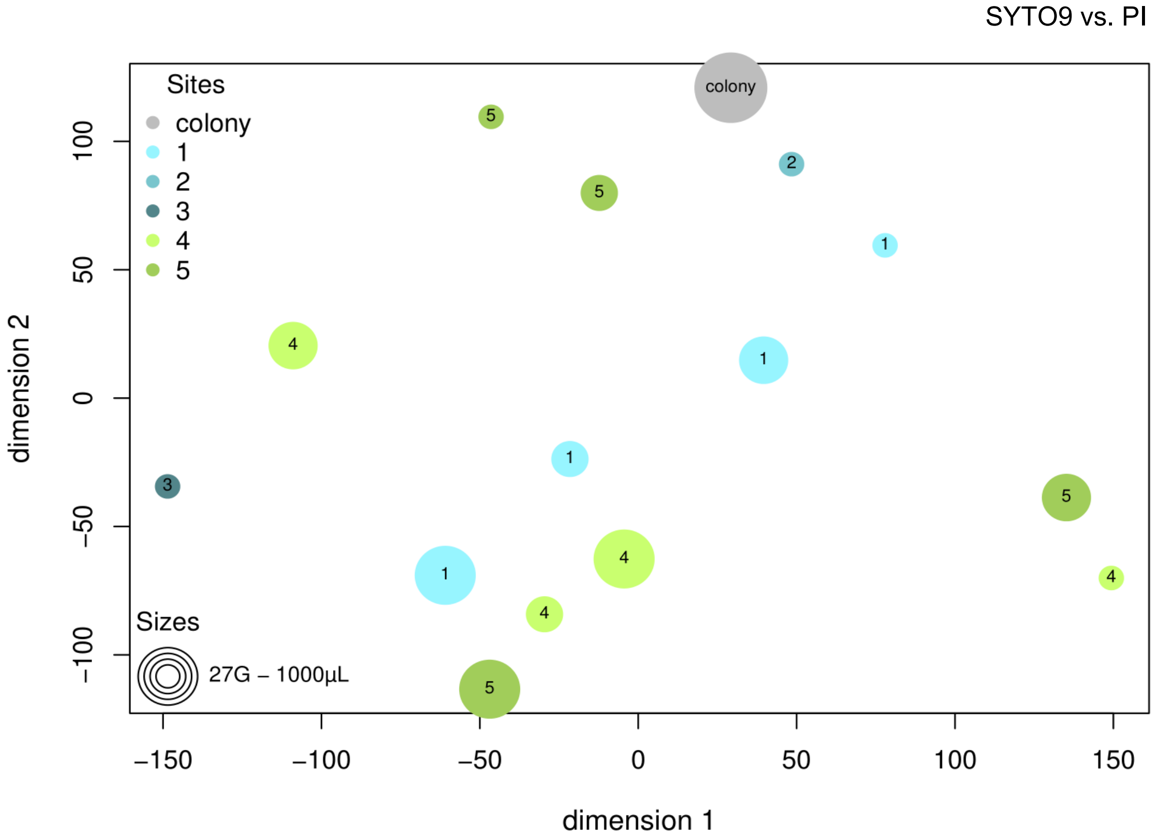


**SI Figure 14**: Similarity comparison of biopsy sites sampled from a 3-day-old *Kocuria rhizophila* colony using different needle or tip sizes. Samples were collected from five distinct positions within the colony (labeled 1 to 5 in respective colors, as detailed in Table 2), and measured by flow cytometry to create cytometric fingerprints (2D plots). The size of the points reflects the dimensions of the sampling tips: 27G needle, as well as 10 µL, 200 µL and 1000 µL pipette tips. Grey point: whole colony. Above: Samples were fixated and stained by the DAPI staining method. Below: Live samples were stained with the SYTO9 and PI staining method. The cell numbers in gates of the 2D plots were transformed by the Box-Cox transformation method and visualized by the R package tSNE.

##### *Stenotrophomonas rhizophila (DSM 14405)*

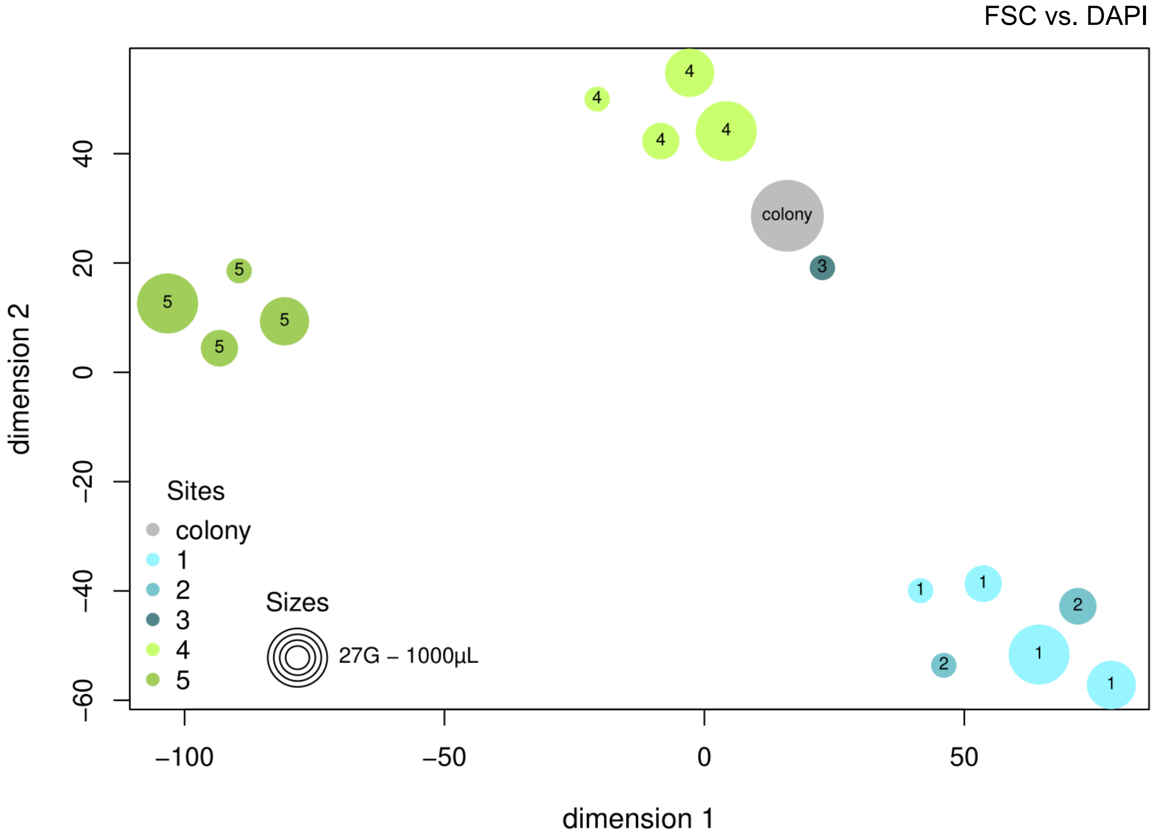


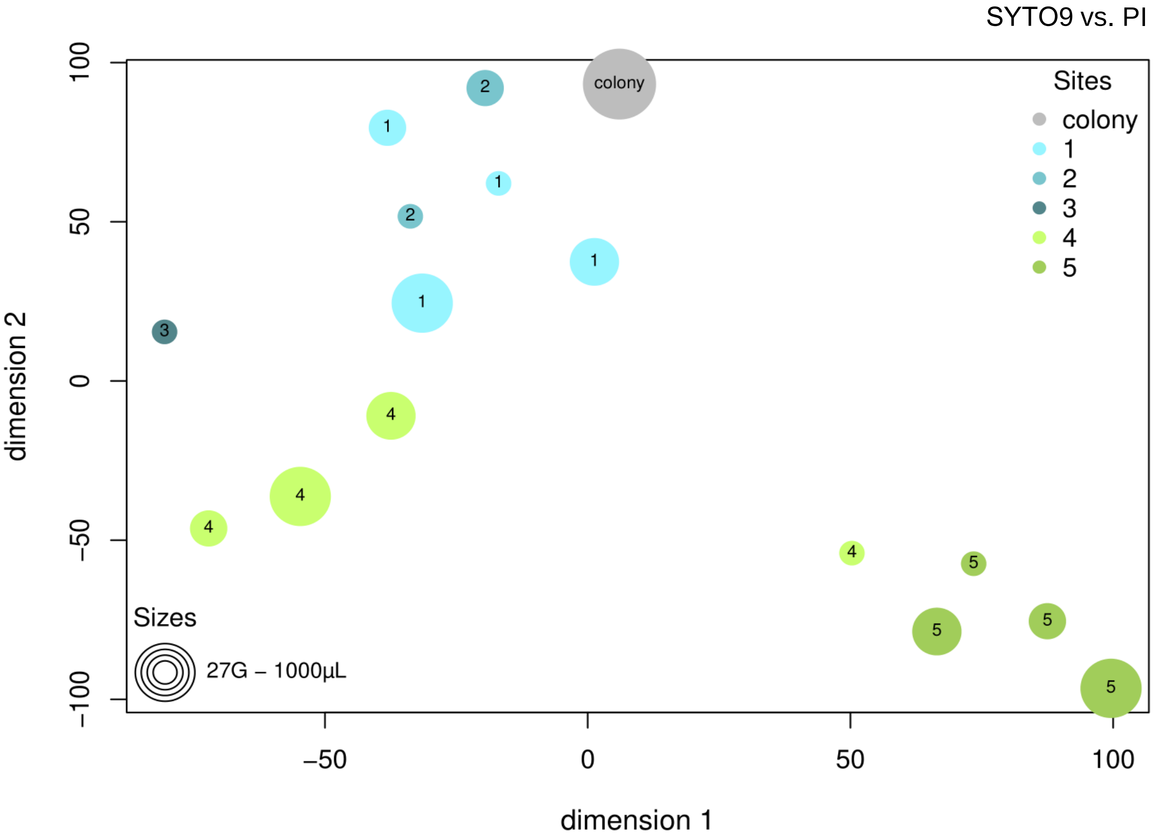


**SI Figure 15**: Similarity comparison of biopsy sites sampled from a 3-day-old *Stenotrophomonas rhizophila* colony using different needle or tip sizes. Samples were collected from five distinct positions within the colony (labeled 1 to 5 in respective colors, as detailed in Table 2), and measured by flow cytometry to create cytometric fingerprints (2D plots). The size of the points reflects the dimensions of the sampling tips: 27G needle, as well as 10 µL, 200 µL and 1000 µL pipette tips. Grey point: whole colony. Above: Samples were fixated and stained by the DAPI staining method. Below: Live samples were stained with the SYTO9 and PI staining method. The cell numbers in gates of the 2D plots were transformed by the Box-Cox transformation method and visualized by the R package tSNE.

***Pseudomonas citronellolis (P3B5)***

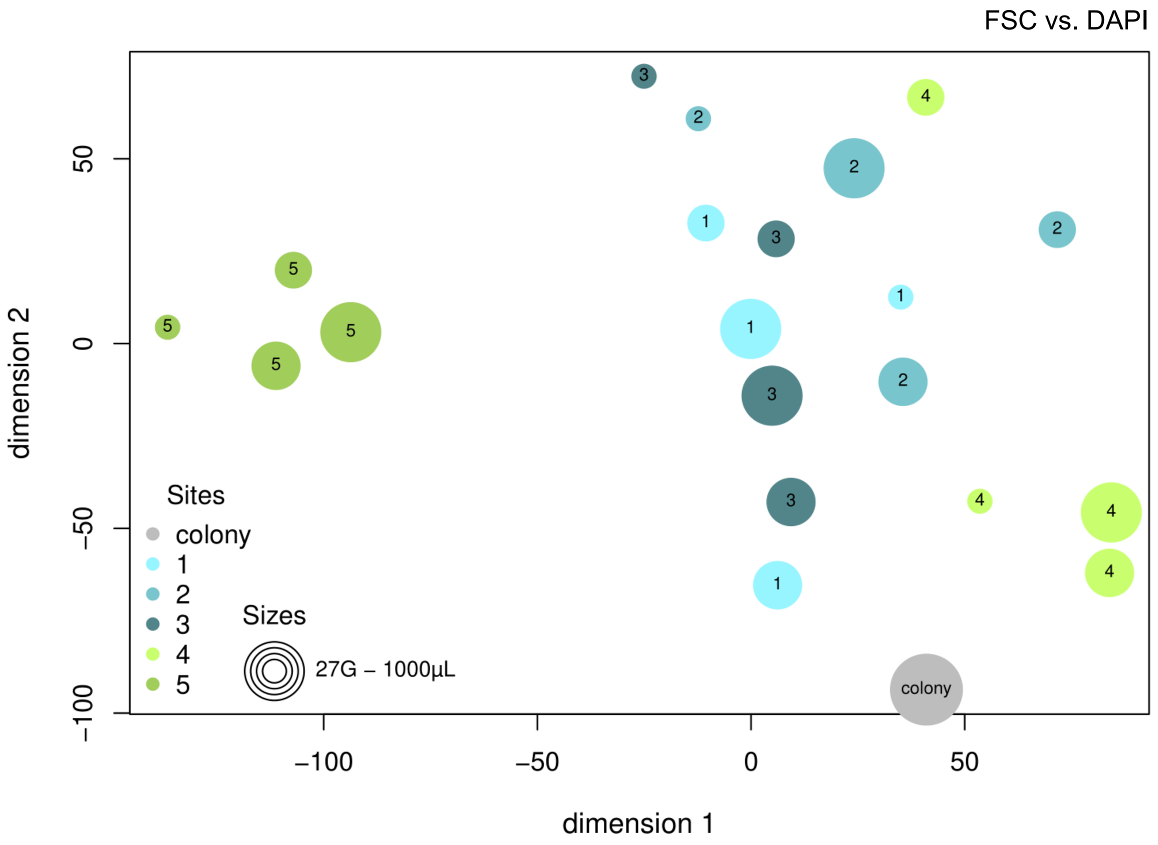


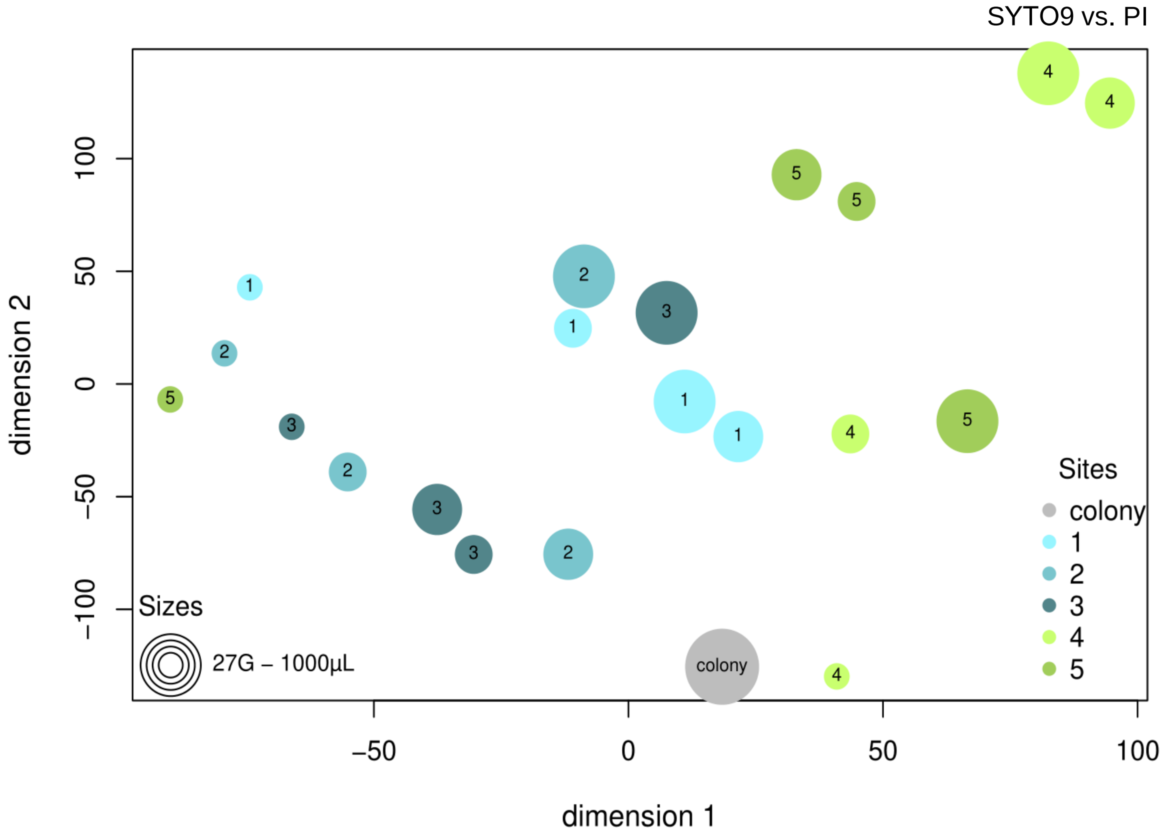


**SI Figure 16:** Similarity comparison of biopsy sites sampled from a 3-day-old *Pseudomonas citronellolis* colony using different needle or tip sizes. Samples were collected from five distinct positions within the colony (labeled 1 to 5 in respective colors, as detailed in Table 2), and measured by flow cytometry to create cytometric fingerprints (2D plots). The size of the points reflects the dimensions of the sampling tips: 27G needle, as well as 10 µL, 200 µL and 1000 µL pipette tips. Grey point: whole colony. Above: Samples were fixated and stained by the DAPI staining method. Below: Live samples were stained with the SYTO9 and PI staining method. The cell numbers in gates of the 2D plots were transformed by the Box-Cox transformation method and visualized by the R package tSNE.

#### *Bacillus subtilis*

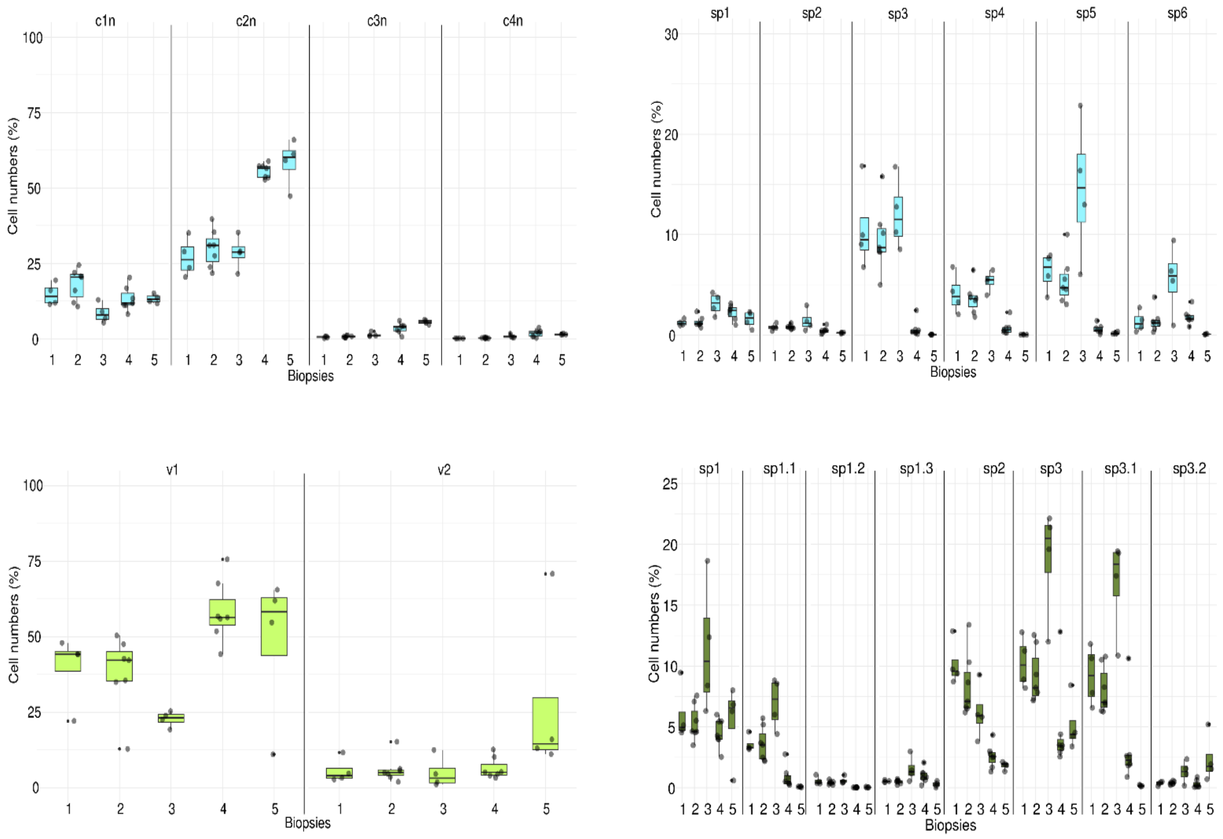


**SI Figure 17**: Proportions of cell types. ABOVE: FSC vs. DAPI analysis. BELOW: SYTO9 vs. PI analysis. LEFT: vegetative cell types. RIGHT: Spore types. To determine the proportion of the various cell types The gate templates in SI Figures 2 and SI Figure 7 were used. Less stained cells were not regarded.

#### *Paenibacillus polymyxa*

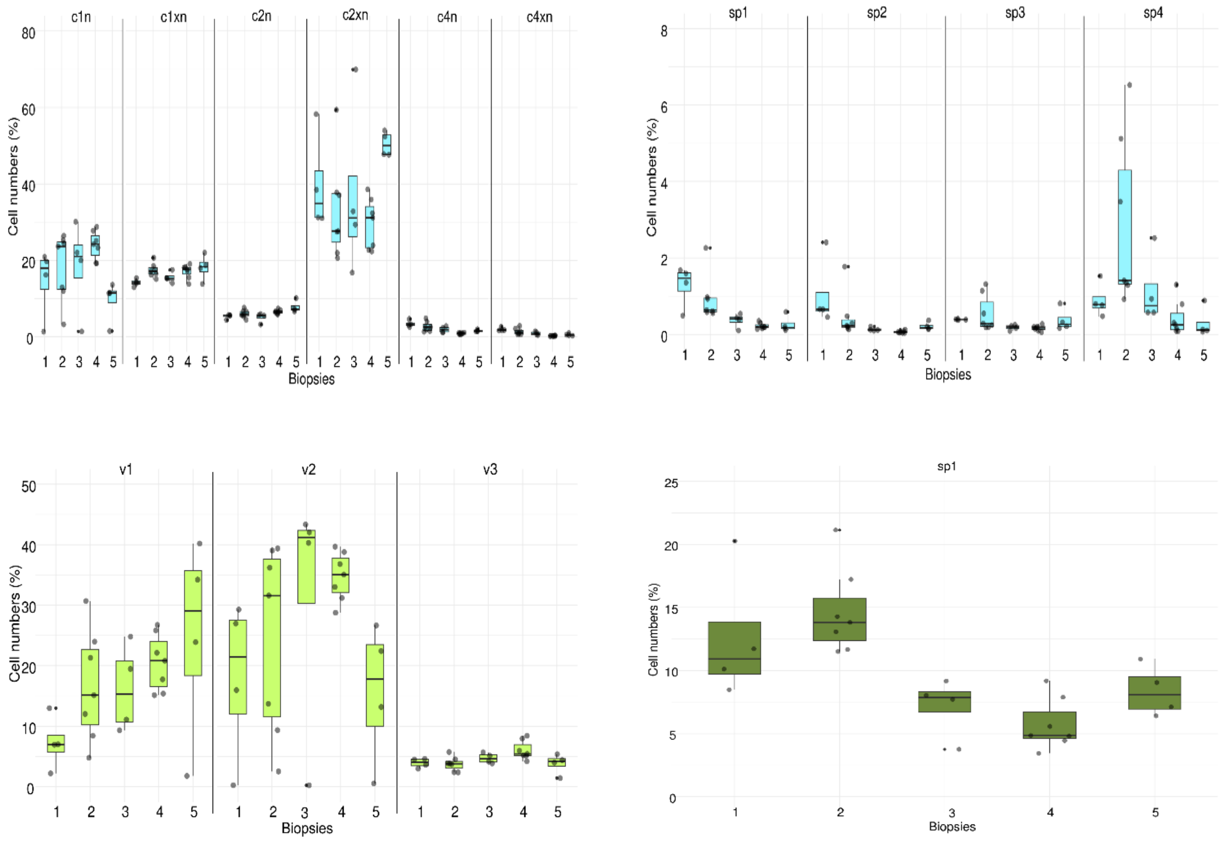


**SI Figure 18**: Proportions of cell types. ABOVE: FSC vs. DAPI analysis. BELOW: SYTO9 vs. PI analysis. LEFT: vegetative cell types. RIGHT: Spore types. To determine the proportion of the various cell types the gate templates in SI Figures 3 and SI Figure 8 were used. Less stained cells were not regarded.

#### *Kocuria rhizophila*

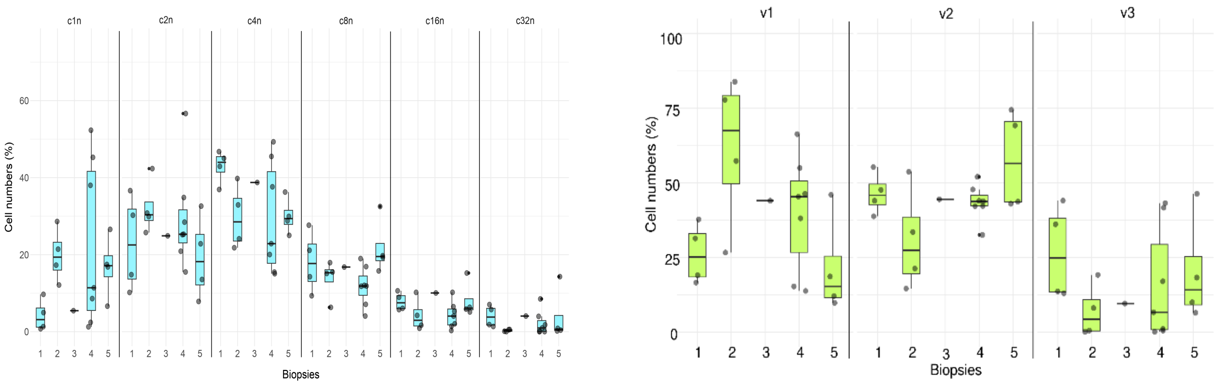


**SI Figure 19**: Proportions of vegetative cell types. LEFT: FSC vs. DAPI analysis. RIGHT: SYTO9 vs. PI analysis. To determine the proportion of the various cell types the gate templates in SI Figures 4 and SI Figure 9 were used.

#### *Stenotrophomonas rhizophila*

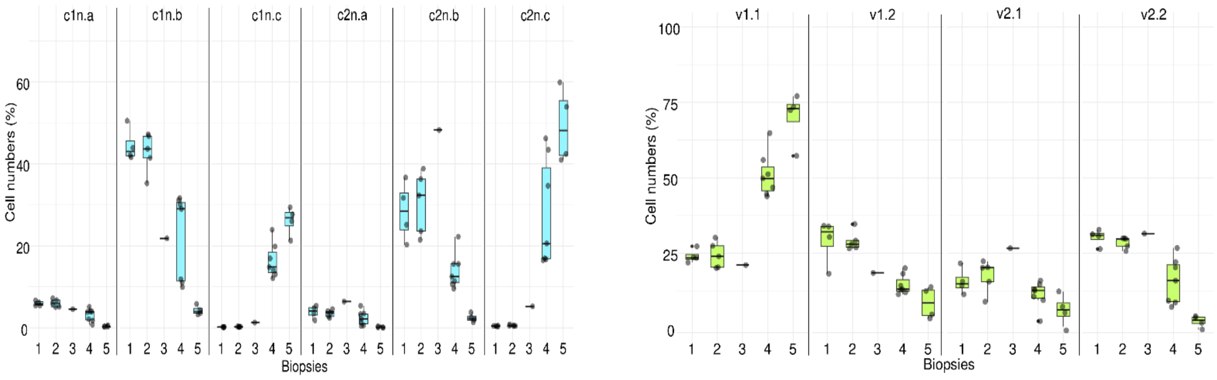


**SI Figure 20**: Proportions of vegetative cell types. LEFT: FSC vs. DAPI analysis. RIGHT: SYTO9 vs. PI analysis. To determine the proportion of the various cell types the gate templates in SI Figures 5 and SI Figure 10 were used. Less stained cells were not regarded.

#### *Pseudomonas citronellolis*

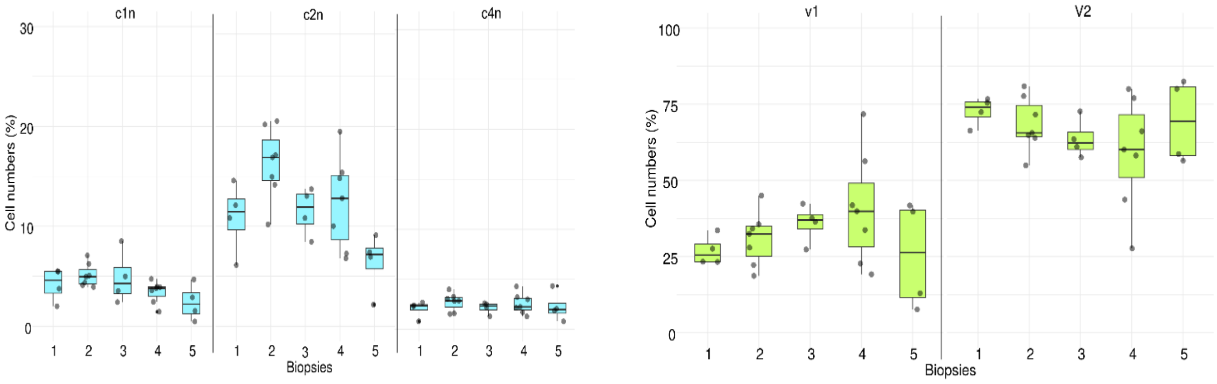


**SI Figure 21**: Proportions of vegetative cell types. LEFT: FSC vs. DAPI analysis. RIGHT: SYTO9 vs. PI analysis. To determine the proportion of the various cell types the gate templates in SI Figures 6 and SI Figure 11 were used. Less stained cells were not regarded.
